# The pore-forming apolipoprotein APOL7C drives phagosomal rupture and antigen cross-presentation by dendritic cells

**DOI:** 10.1101/2023.08.11.553042

**Authors:** Gerone A. Gonzales, Song Huang, Jahanara Rajwani, Liam Wilkinson, Jenny A. Nguyen, Cassandra M. Wood, Irene Dinh, Melanie Moore, Eymi Cedeño, Saif Sikdar, Neil McKenna, Vincent Ebacher, Nicole L. Rosin, Matheus B. Carneiro, Bas Surewaard, Nathan C. Peters, Jeff Biernaskie, Douglas J. Mahoney, Robin M. Yates, Johnathan Canton

## Abstract

Type I conventional dendritic cells (cDC1s) are essential for the generation of protective cytotoxic T lymphocyte (CTL) responses against many types of viruses and tumours. They do so by internalizing antigens from virally infected or tumour cells and presenting them to CD8^+^ T cells in a process known as cross-presentation (XP). Despite the obvious biological importance of XP, the molecular mechanism(s) driving this process remain unclear. Here, we show that a cDC-specific pore-forming protein called apolipoprotein 7C (APOL7C) is upregulated in response to innate immune stimuli and is recruited to phagosomes. Strikingly, the association of APOL7C with phagosomes leads to phagosomal rupture, which in turn allows for the escape of engulfed protein antigens to the cytosol where they can be processed via the endogenous major histocompatibility complex (MHC) class I antigen processing pathway. We show that APOL7C recruitment to phagosomes is voltage-dependent and occurs in response to NADPH oxidase-induced depolarization of the phagosomal membrane. Our data indicate the presence of dedicated pore-forming apolipoproteins that mediate the delivery of phagocytosed proteins to the cytosol of activated cDC1s to facilitate MHC class I presentation of exogenous antigen and to regulate adaptive immunity.

## Introduction

Antigen-presenting cells (APCs), such as macrophages and dendritic cells, can present exogenous antigens on major histocompatibility complex class I (MHC-I) to naïve CD8^+^ T cells to elicit cytotoxic T lymphocyte (CTL) responses. This process, referred to as cross-presentation (XP), is essential to the clearance of many types of viruses and tumours^1–8^. While several APCs are capable of XP, experiments using *Batf3-*deficient (*Batf3^-/-^*) or *Irf8* + 32*^-/-^* mice, which both lack cDC1s, have demonstrated that cDC1s play a seemingly nonredundant role in the generation of CTLs^8–10^. Indeed, cDC1s harbor unique specializations that facilitate their role in XP. For example, they express the receptor DNGR-1 (also known as CLEC9A) which facilitates the XP of antigens extracted from dead or damaged cells^5,7,11–14^. Similarly, cDC1s uniquely express high levels of WDFY4, a protein that is required for XP by an unknown mechanism^1^. Importantly, the molecular mechanisms by which XP is achieved likely vary depending on the environmental context and the nature of the antigen.

Two pathways for XP are believed to exist. In the first, known as the “vacuolar” pathway, proteins are processed by endosomal proteases to liberate peptides which are loaded onto MHC-I all within endocytic organelles^15–17^. The second is known as the “cytosolic” or the “phagosome-to-cytosol (P2C)” pathway^15–17^. In this pathway, endocytosed or phagocytosed proteins are released into the cytosol where they are processed into peptides by the proteasome. The peptides are then loaded onto MHC-I in the ER for XP. The mechanism by which P2C occurs is ill-defined and multiple hypotheses exist^19^. One hypothesis, the “indigestion model”, proposes that under certain conditions the membrane integrity of phagosomes is lost allowing for the release of engulfed proteins into the cytosol^20,21^. Recently, evidence in support of the “indigestion model” has emerged. For example, DNGR-1 has been found to signal via SYK and the NADPH oxidase for phagosomal rupture, which in turn releases phagocytosed proteins into the cytosol^11,22^. Also, a pore-forming protein present in the lumen of endocytic organelles in APCs called perforin-2 has been found to instigate the formation of non-selective pores in endocytic organelle membranes, which allows for the escape of endocytosed proteins to the cytosol^23^. These exciting new findings have led to entirely new questions. For example, do pore-forming proteins such as perforin-2 contribute to DNGR-1 dependent phagosomal rupture, what signals regulate rupture or pore formation in the endocytic pathway of APCs, does it occur on all endocytic organelles, and are there other, yet undefined, molecular drivers of this process?

In this study, we present new evidence in support of the “indigestion model” for P2C. We show that innate immune stimuli, including agonists of Toll-like receptor 3 (TLR3) which is highly expressed by cDC1s, lead to the expression of apolipoprotein L 7C (APOL7C), a cytosolic apolipoprotein L containing a putative colicin-like pore-forming domain. We find that APOL7C inserts into phagosomes in an NADPH oxidase-dependent manner, leading to pore formation and causing phagosomal rupture and the release of phagocytosed proteins to the cytosol for XP. Our data indicate the presence of dedicated cDC-specific pore-forming proteins that insert into phagosomes in a regulated manner to drive XP.

## Results

### Poly(I:C) leads to a greater incidence of phagosomal membrane rupture in cDC1s

Previous studies have shown that XP is a tuneable process^24–26^ that can be enhanced by innate immune stimuli^24,25^ and both the TLR4 agonist LPS and the TLR9 agonist CpG DNA can result in enhanced damage to endosomes in dendritic cells^27,28^. To investigate the effect of cDC1 activation on phagosomal rupture in cDC1s, we employed the murine cDC1 cell line MuTuDC1940 (henceforth called MuTuDCs). MuTuDCs are frequently used to study XP, have a similar proteomic profile to primary cDC1s *in vivo,* and have shown evidence of phagosomal rupture^11,23,28–34^. We first treated MuTuDCs with poly(I:C), which can trigger TLR3 that is particularly abundant on cDC1s and favours enhanced cross-priming^31,35–42^. As expected, treatment of MuTuDCs with poly(I:C) led to enhanced XP of ovalbumin that was covalently coupled to beads (OVA-beads) and, to a lesser extent, also increased the direct presentation of soluble OVA peptide (SIINFEKL) (**Figure 1A-B**). The number of phagocytosed OVA-beads was not affected by poly(I:C) treatment (**Figure 1C-D**). Using these conditions, we next assessed whether poly(I:C)-dependent activation of MuTuDCs had any effect on phagosomal rupture.

**Figure 1.**
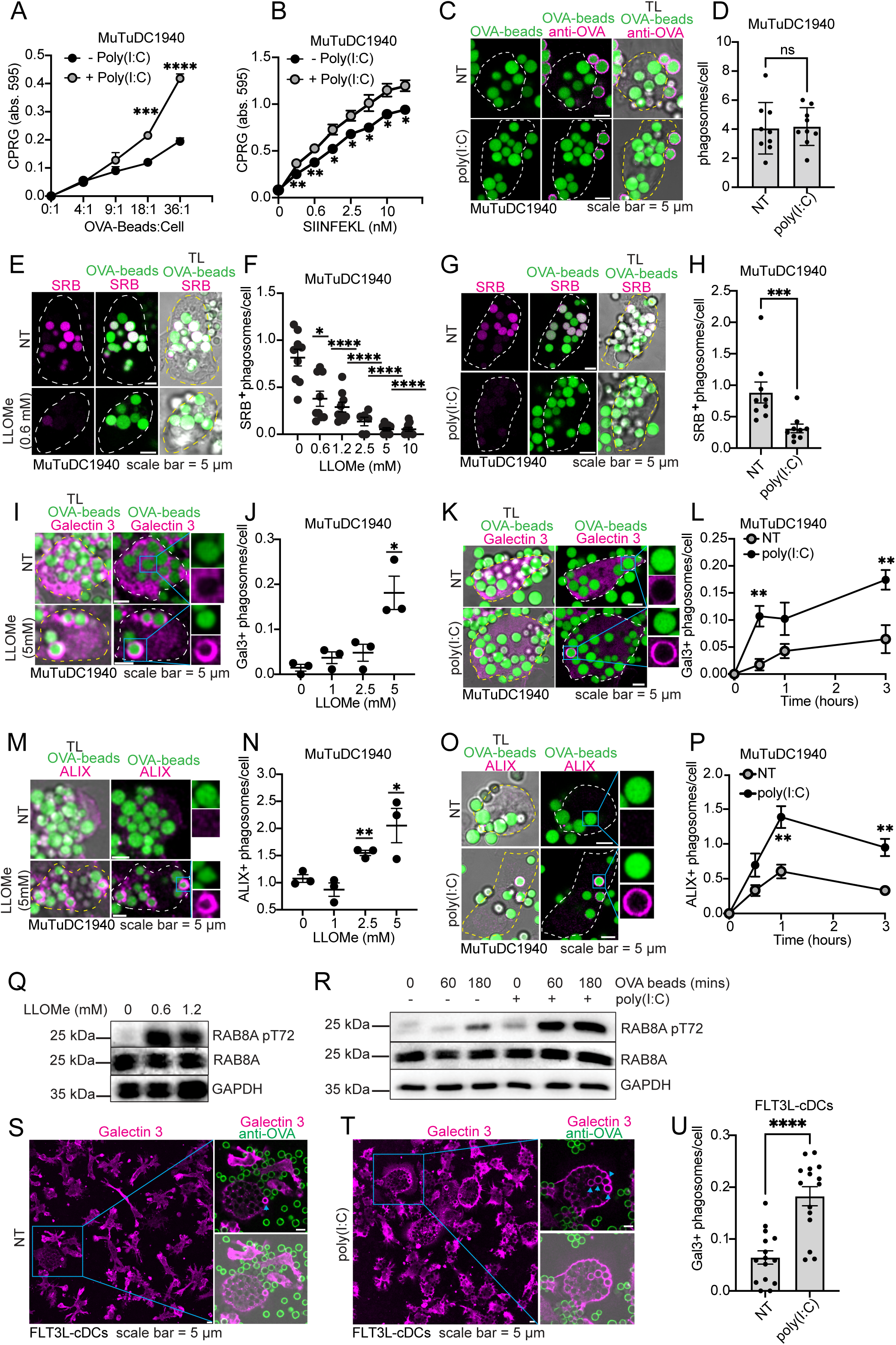
Poly(I:C) treatment increases the incidence of phagosomal rupture in MuTuDCs and FLT3L-cDCs. **(A-B)** MuTuDCs were either left untreated or treated with poly(I:C) at 5 μg/ml overnight. After a 4hr incubation with OVA-beads **(A)** or SIINFEKL **(B)**, the cells were fixed and incubated overnight with B3Z cells. B3Z cells were lysed in a CPRG containing buffer. Plotted as mean ± s.d. of experimental triplicates and is representative of 10 independent experiments (*n* = 10). Significance determined using student’s *t* test for individual dilutions. **(C)** MuTuDCs were incubated with OVA-beads for 4 hours before they were fixed and stained for uninternalized beads with an anti-OVA antibody. Images were acquired by confocal microscopy. Scale bar = 5 μm. **(D)** The number of internalized OVA-beads were counted from the micrographs shown in **(C)**. Plotted as mean ± s.d. Each dot represents a field-of-view containing 5-20 cells (*n* = 3 independent experiments). Significance was determined using a Kolmogorov-Smirnov test. **(E-F)** MuTuDCs were incubated with OVA-beads and SRB for 1hr. SRB was washed off and replaced with medium containing LLOMe at the indicated concentration. Cells were imaged live by confocal microscopy. Scale bar = 5 μm. The number of SRB^+^ phagosomes per cell is shown. Plotted as mean ± s.e.m. Each dot represents a field of view containing 5-20 cells, compiled from 3 independent experiments (*n* = 3). Significance determined using student’s *t* test against untreated sample. **(G-H)** MuTuDCs were treated or not with poly(I:C) (5 μg/ml) overnight. SRB leakage was assessed as in **(E-F)**. Scale bar = 5 μm. The number of SRB^+^ phagosomes per cell is shown. Plotted as mean ± s.e.m. Each dot represents a field of view containing 5-20 cells, compiled from 3 independent experiments (*n* = 3). Significance determined using Kolmogorov-Smirnov test. **(I-J)** MuTuDCs were incubated with OVA-beads for 1hr. The medium was replaced with medium containing LLOMe at the indicated concentration. Cells were stained for Galectin 3 and imaged live by confocal microscopy. Scale bar = 5 μm. The number of Galectin 3 positive phagosomes per cell is shown. Plotted as mean ± s.e.m. of experimental triplicates, each dot represents a field of view containing 5-20 cells, representative of 2 independent experiments (*n* = 2). Significance determined using student’s *t* test against untreated sample. **(K-L)** MuTuDCs were treated or not with poly(I:C) (5 μg/ml) overnight. Galectin 3 recruitment was assessed as in **(I-J)**. Scale bar = 5 μm. The number of Galectin 3 positive phagosomes per cell is shown. Plotted as mean ± s.e.m. of experimental triplicates and is representative of 3 independent experiments (*n* = 3). Each dot represents a field of view containing 5-20 cells. Significance determined using student’s *t* test. **(M-N)** MuTuDCs were incubated with OVA-beads for 1hr. The medium was replaced with medium containing LLOMe at the indicated concentration. Cells were stained for ALIX and imaged live by confocal microscopy. Scale bar = 5 μm. The number of ALIX positive phagosomes per cell is shown. Plotted as mean ± s.e.m. of experimental triplicates, each dot represents a field of view containing 5-20 cells, representative of 2 independent experiments (*n* = 2). **(O-P)** MuTuDCs were treated or not with poly(I:C) (5 μg/ml) overnight. ALIX recruitment was assessed as in **(M-N)**. Scale bar = 5 μm. The number of ALIX positive phagosomes per cell is shown. Plotted as mean ± s.e.m. of experimental triplicates and is representative of 3 independent experiments (*n* = 3). Each dot represents a field of view containing 5-20 cells. Significance determined using student’s *t* test. **(Q)** Western blot for phosphorylated RAB8A, RAB8A and GAPDH. MuTuDCs treated with LLOMe for 1hr at the indicated concentration. **(R)** Western blot for phosphorylated RAB8A, RAB8A and GAPDH. MuTuDCs were treated with poly(I:C) (5 μg/ml) overnight and then challenged with OVA-beads for the indicated time. **(S-U)** FLT3L-cDCs were treated or not with poly(I:C) (5 μg/ml) overnight. Galectin 3 recruitment was assessed as in **(I-J)**. Scale bar = 5 μm. The number of Galectin 3 positive phagosomes per cell is shown. Plotted as mean ± s.e.m. Each dot represents a field of view containing 5-20 cells and plot is compiled from 3 independent experiments (*n* = 3). Significance determined using student’s *t* test. n.s. = no significance; \**P* ≤ 0.05; \*\**P* ≤ 0.01; \*\*\**P* ≤ 0.001; \*\*\*\**P* ≤ 0.0001.

We established several independent assays to assess the incidence of phagosomal rupture in MuTuDCs upon poly(I:C) activation. In the first, OVA-beads were covalently labeled with a fluorophore (Alexa Fluor 488; AF488) and incubated with MuTuDCs in the presence of the fluid-phase, membrane-impermeant dye sulforhodamine B (SRB). The uptake of the OVA-beads via phagocytosis results in the simultaneous loading of phagosomes with SRB. Unlike latex beads, the 3 μm silica beads used here are porous allowing for the entry of fluorescent molecules like SRB^43^; as a result, the fluid-phase SRB is visible throughout the phagosome. Upon phagosomal rupture, fluid-phase SRB escapes into the cytosol, but covalently attached AF488 remains within the phagosome. Using the phagolysosome-damaging compound L-leucyl-L-leucine methyl ester (LLOMe), we confirmed that phagosomal rupture results in the progressive loss of SRB, but not the AF488, from phagosomes (**Figure 1E-F**). Next, we assessed the effect of poly(I:C) activation on the loss of SRB from phagosomes. Interestingly, MuTuDCs treated with poly(I:C) displayed significantly fewer SRB-positive phagosomes relative to untreated cells (**Figure 1G-H**) suggestive of phagosomal rupture. Therefore, we next looked at the incidence of established phagosomal damage biomarkers in poly(I:C) treated cells.

Damage to endocytic organelle or phagosome membranes initiates cellular membrane repair processes characterized by the rapid phosphorylation of RAB8A by the kinase LRRK2 which subsequently recruits components of the endosomal sorting complex required for transport (ESCRT)-III including CHMP4B and ALIX^44–50^. In the case of extensive damage, members of the galectin family of proteins are also recruited to phagosomes to facilitate removal of the damaged endocytic organelles by autophagy^46–48,50^. As a result, recruitment of components of these membrane repair/disposal pathways to endocytic organelles or phagosomes are frequently used as indirect biomarkers of membrane damage^11,28,34,46,49,51–54^. Indeed, we found that treatment of MuTuDCs with the phagosome-damaging compound LLOMe resulted in a dose-dependent increase in the number of galectin 3- and ALIX-positive phagosomes (**Figure 1I-J and 1M-N**). Similarly, RAB8A was phosphorylated in MuTuDCs harboring OVA-beads upon LLOMe treatment (**Figure 1Q**). Using these markers of endocytic organelle/phagosome damage, we assessed the incidence of damaged phagosomes in poly(I:C) treated cells. Interestingly, we found a significant increase in the number of galectin 3- and ALIX-positive phagosomes in poly(I:C)-treated cells relative to untreated cells at various time-points post-uptake (**Figure 1K-L and 1O-P**). Likewise, poly(I:C) treatment significantly enhanced RAB8A phosphorylation at various timepoints after the addition of OVA-beads (**Figure 1R**). Next, we assessed whether similar findings could be observed in FLT3L-driven cultures of bone marrow cells (henceforth referred to as FLT3L-cDCs), which generate primary cDC1 and cDC2 subsets (**Supplementary Figure 1A-B**). As with the MuTuDCs, treatment of FLT3L-cDCs with poly(I:C) resulted in a significant increase in the number of galectin 3-positive phagosomes per cell (**Figure 1S-U**). Taken together, our data suggest that poly(I:C) treatment results in a greater incidence of phagosome rupture in MuTuDCs and primary FLT3L-cDCs.

### Poly(I:C) increases expression of the pore-forming apolipoprotein APOL7C which is recruited to phagosomes

To gain insight into the molecular pathways that drive phagosomal rupture upon poly(I:C) treatment of cDC1s, we treated MuTuDCs with or without poly(I:C) and performed bulk RNA sequencing (RNAseq). Among others, we found a significant increase in a group of proteins known as the apolipoprotein L (APOL) family (**Figure 2A**). Intriguingly, APOL proteins harbor putative colicin-like pore-forming domains that are known to induce damage to biological membranes, including endocytic organelles^55–58^. While several members of the family were upregulated upon poly(I:C) treatment, apolipoprotein L 7C (APOL7C) demonstrated a greater than 20-fold increase in expression (**Figure 2B-C**). We confirmed that this induction also occurred in splenic cDCs *in vivo* by injecting mice with poly(I:C) via the tail vein, isolating splenocytes, enriching for cDCs and performing qRT-PCR. Here again we found a significant increase in *Apol7c* expression (**Figure 2D**).

**Figure 2.**
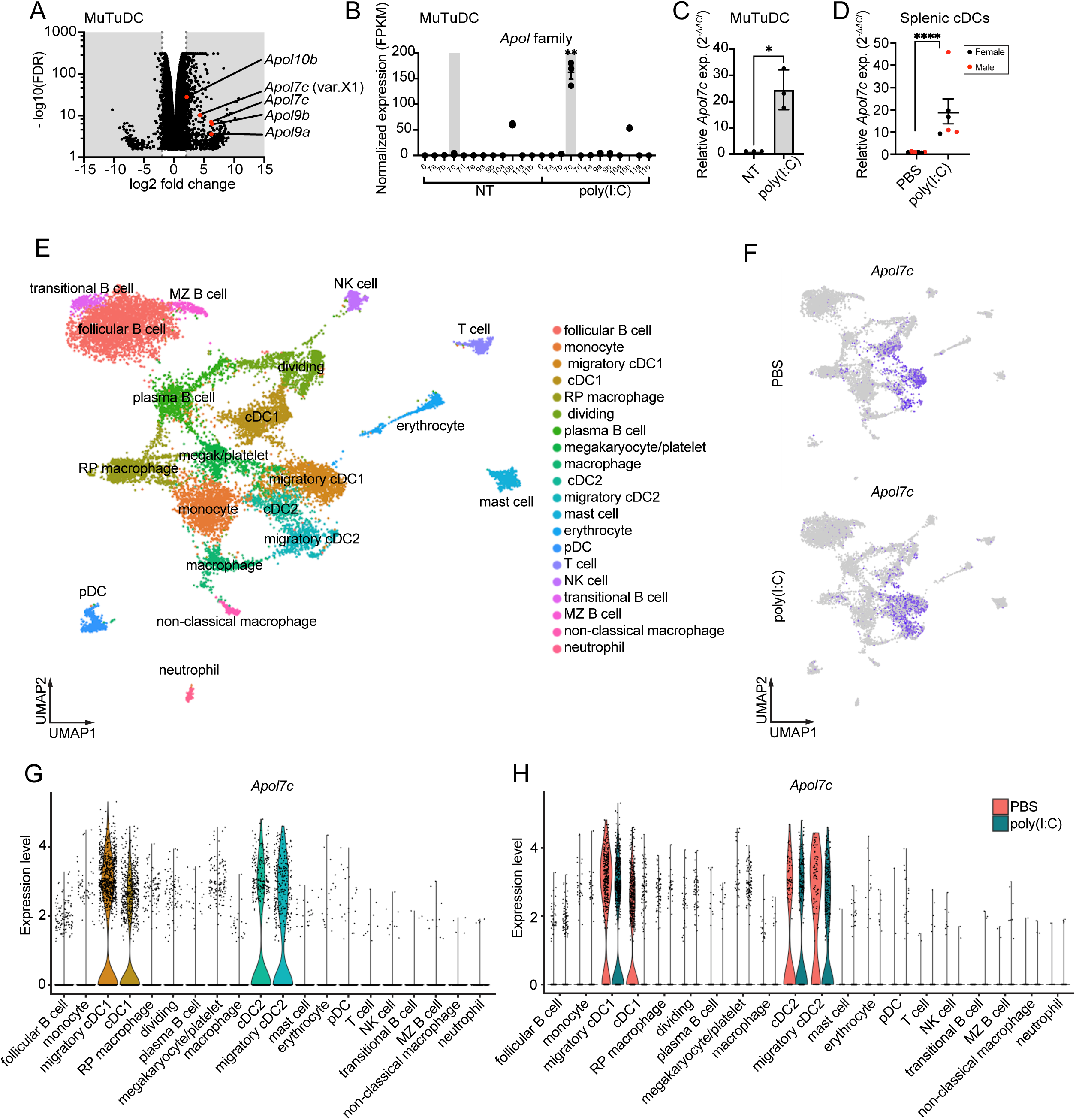
*Apol7c* expression is enhanced by poly(I:C) and is unique to cDCs. **(A)** Volcano plot derived from bulk RNAseq of MuTuDCs treated with or without poly(I:C) (5 μg/ml) overnight. Compiled from 3 independent experiments (*n* = 3). **(B)** Normalized expression of all murine APOL family members from bulk RNAseq experiment described in **(A)**. Each dot represents the mean ± s.e.m. from 3 independent experiments (*n* = 3). Significance determined using student’s *t* test. **(C)** MuTuDCs were treated with or without poly(I:C) (5 μg/ml) overnight and *Apol7c* expression was assessed by RT-qPCR. Each dot represents the mean ± s.d. from 3 independent experiments (*n* = 3). Significance determined using student’s *t* test. **(D)** Wild-type C57BL/6 mice were treated with poly(I:C) (50 μg/mouse) via tail vein. 24hr later the spleens were harvested and RNA was prepared from a single cell suspension of spleen cells. *Apol7c* expression was assessed by RT-qPCR. Plotted as mean ± s.e.m. (*n* = 3 mice per group). Significance determined using student’s *t* test. **(E)** Uniform manifold approximation and projection (UMAP) of pooled murine spleen CD45^+^ and pan-DC cells (n = 17,027 cells). **(F)** UMAPs showing expression of *Apol7c* by color (gray, low expression; blue, high expression). **(G-H)** Violin plots showing the expression of *Apol7c* by splenic cell populations in mice that were untreated **(G)** or treated with poly(I:C) **(H)** from scRNAseq data. n.s. = no significance; \**P* ≤ 0.05; \*\**P* ≤ 0.01; \*\*\*\**P* ≤ 0.0001.

We next assessed the immune cell-specificity of *Apol7c* expression by single-cell RNA-sequencing (scRNAseq). To this end, we treated mice with poly(I:C) or PBS by tail vein injection and after 24 hours processed splenocytes for scRNAseq. An unsupervised uniform manifold approximation and projection (UMAP) was performed and the annotation of cell clusters was based on the expression of curated and data-driven genes as described by Bosteels *et al*^59^, Rawat *et al*^60^, Brown *et al*^61^, and Sheu *et al*^62^. A total of 19 clusters were identified where clusters 2 (*Ccr7, Laptm4b, Ccl5, Fscn1, Tmem123, Crip1, AW112010, Basp1*)^59,60^, 3 (*Clec9a, Cd36, Tlr3, Irf8, Cst3, Cadm1, Rab7b,*)^59,60^, 9 (*S100a4, Relb, Tmem176a, Tmem176b, Tbc1d4,*)^59,61^ and 10 (*Ccr7, Relb, Tmem176a, Il4i1, Tmem123, Cd83, Socs2*)^59,60^ represented migratory cDC1s^59,60^, non-migratory cDC1s^59,60^, non-migratory cDC2s^59,61^ and migratory cDC2s^59,60^, respectively (**Figure 2E, Supplementary Figure 2** and **Supplementary Table 1**). Interestingly, we found that *Apol7c* expression was restricted to cDCs, with the highest expression in migratory cDC1s (**Figure 2E-H**). In addition to migratory cDC1s, *Apol7c* expression was also found in non-migratory cDC1s, migratory cDC2s and non-migratory cDC2s (**Figure 2E-H**). Poly(I:C) treatment did not change this pattern, as *Apol7c* expression remained restricted to cDCs (**Figure 2H**).

We next turned our attention to the subcellular distribution of APOL7C. For this we generated a stable MuTuDC cell line expressing a doxycycline(dox)-inducible APOL7C::mCherry fusion protein (henceforth referred to as MuTuDC.APOL7C::mCherry cells). We found APOL7C::mCherry to be largely cytosolic in otherwise untreated cells. Remarkably, upon phagocytosis of OVA beads, APOL7C::mCherry redistributed to phagosomes (**Figure 3A-B**). Interestingly, we saw similar phagosomal recruitment of APOL7C::mCherry to phagosomes in another phagocyte cell line Raw 264.7, suggesting that while APOL7C is selectively expressed by cDCs, its ability to be recruited to phagosomes can be reconstituted in other cells (**Figure 3E**). Phagosomal recruitment of APOL7C::mCherry was not limited to OVA-bead-containing phagosomes, as we observed similar recruitment to zymosan-, IgG-opsonized sheep red blood cell (IgG-sRBC)-, *Leishmania major*- and *Staphylococcus aureus*-containing phagosomes (**Supplementary Figure 3A-D**). Recruitment of APOL7C::mCherry to phagosomes appeared to peak several hours after phagocytosis suggesting that it was recruited to late phagosomes that have likely already fused with the lysosomal compartment (**Figure 3B**). To confirm this, we transiently transfected Raw264.7 cells stably expressing H-2K^b^ and dox-inducible APOL7C::mCherry (henceforth termed RawKb.APOL7C::mCherry) with p40PX::GFP which is a biosensor for phosphatidylinositol-(3)-phosphate [PtdIns(3)P]. PtdIns(3)P is only present on newly formed or early phagosomes. We found no phagosomes that were positive for both APOL7C::mCherry and p40PX::GFP (**Supplementary Figure 3E-G**). Next, we stained cells expressing APOL7C::mCherry for a marker of late phagosomes, LAMP1. We found that nearly all phagosomes that were positive for APOL7C::mCherry were also positive for LAMP1 (**Supplementary Figure 3H-J**). Together, these data demonstrate that APOL7C::mCherry accumulates on late phagosomes that have already fused with lysosomes (i.e., phagolysosomes). Given that APOL7C harbors a putative pore-forming domain and that other APOL family members have been shown to permeabilize biological membranes, we next assessed whether APOL7C contributes to phagosomal rupture.

**Figure 3.**
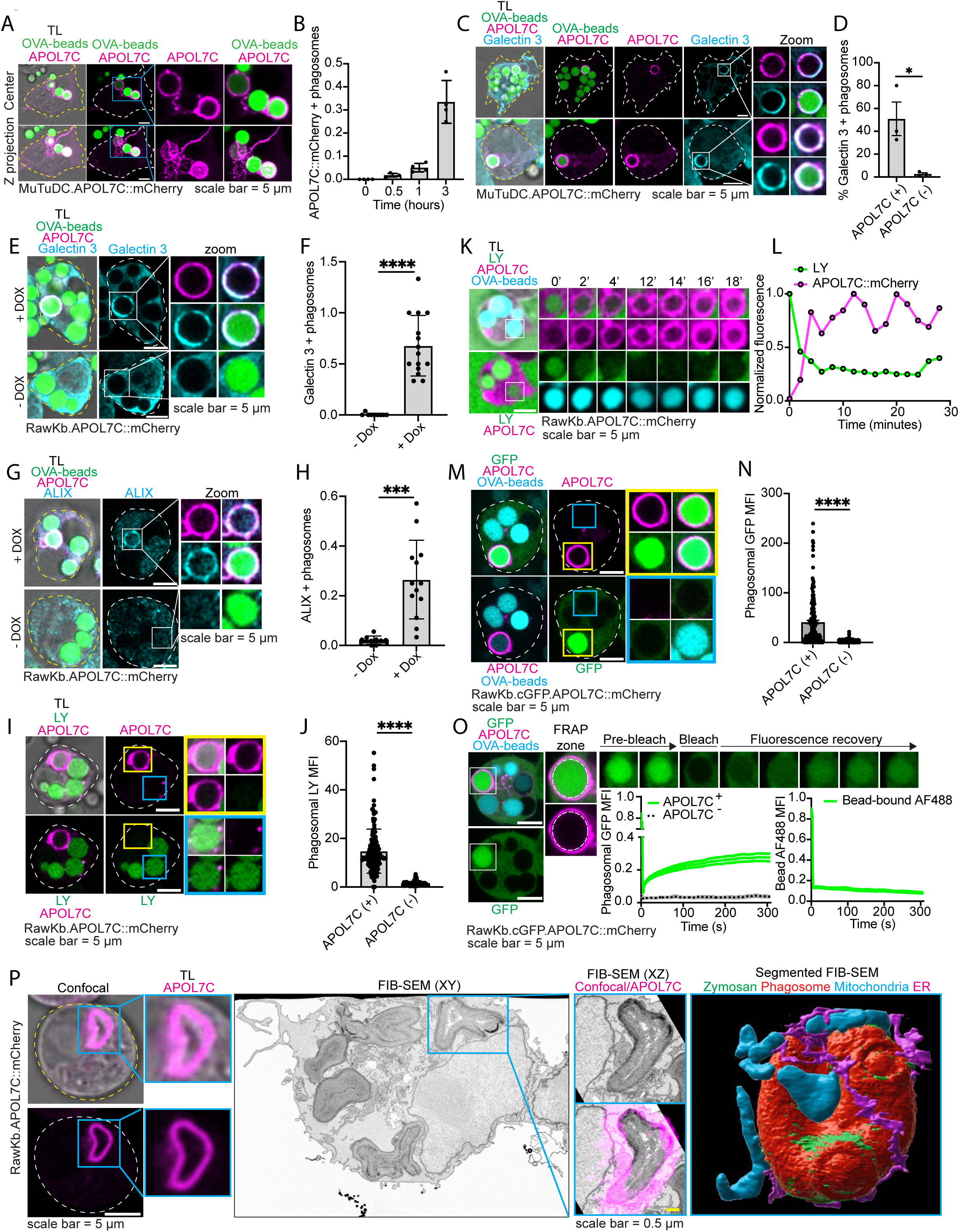
APOL7C is recruited to phagosomes and instigates phagosomal rupture. **(A-B)** MuTuDC.APOL7C::mCherry cells were treated with doxycycline (DOX) overnight and then incubated with OVA-beads for the indicated time before fixation and confocal imaging. Scale bar = 5 μm. Plotted as mean ± s.e.m. Each dot represents a field of view containing 5-20 cells from 4 independent experiments (*n* = 4). **(C-D)** MuTuDC.APOL7C::mCherry cells were treated with doxycycline (DOX) overnight and then incubated with OVA-beads for 4hr before fixation, staining for galectin 3 and confocal imaging. Scale bar = 5 μm. Plotted as mean ± s.e.m. of the percentage of APOL7C^+^ or APOL7C^-^ phagosomes that were positive for Galectin 3 from 5 fields of view containing 5-20 cells each. Each dot is an independent experiment (*n* = 3). A total of 160 phagosomes were assessed. Significance determined using student’s *t* test. **(E-F)** RawKb.APOL7C::mCherry cells were treated with or without doxycycline (DOX) overnight and then incubated with OVA-beads for 4hr before fixation, staining for galectin 3 and confocal imaging. Scale bar = 5 μm. Plotted as mean ± s.d. of the number of APOL7C^+^ phagosomes/cell from 9-15 fields of view containing 5-20 cells each. Each dot is representative of a field of view, compiled from 3 independent experiments (*n* = 3). Significance determined using student’s *t* test. **(G-H)** RawKb.APOL7C::mCherry cells were treated with doxycycline (DOX) overnight and then incubated with OVA-beads for 4hr before fixation, staining for ALIX and confocal imaging. Scale bar = 5 μm. Plotted as mean ± s.d. of the number of APOL7C^+^ phagosomes/cell that were positive for ALIX from 9-15 fields of view containing 5-20 cells each. Each dot is representative of a field of view, compiled from 3 independent experiments (*n* = 3). Significance determined using student’s *t* test. **(I-J)** RawKb.APOL7C::mCherry cells were treated with doxycycline (DOX) overnight and then incubated with OVA-beads and Lucifer Yellow (LY) for 1hr. The LY was washed away and the cells were incubated for an additional 3hr before live cell confocal imaging. Scale bar = 5 μm. Plotted as mean ± s.d. of the mean fluorescence intensity (MFI) of LY in either APOL7C^+^ or APOL7C^-^ phagosomes. Each dot is representative of the MFI of a single phagosome, compiled from 3 independent experiments (*n* = 3). Significance determined using a Kolmogorov-Smirnov test. **(K-L)** RawKb.APOL7C::mCherry cells were treated as in **(I)** and then imaged live on the confocal microscope. Plotted is the MFI of LY and APOL7C::mCherry for the indicated phagosome. **(M-N)** RawKb.cGFP.APOL7C::mCherry cells were treated with doxycycline (DOX) overnight and then imaged live by confocal microscopy. Scale bar = 5 μm. Data is plotted as the mean ± s.d. of the MFI of GFP in the lumen of APOL7C^+^ and APOL7C^-^ phagosomes. Each dot represents a single phagosome, compiled for 3 independent experiments (*n* = 3). Significance determined using student’s *t* test. **(O)** RawKb.cGFP.APOL7C::mCherry cells were treated with doxycycline (DOX) overnight and then imaged live by confocal microscopy. Phagosomes were photobleached and the fluorescence recovery was recorded for the time indicated (*n* = 15 phagosomes for APOL7C^+^, 11 phagosomes for APOL7C^-^, and 11 phagosomes for AF488). **(P)** RawKb.APOL7C::mCherry cells were treated with doxycycline (DOX) overnight and then with zymosan for 3hr. Cells were then imaged by live cell confocal microcospy and processed for FIB-SEM. Scale bars = 5 μm for confocal and 0.5 μm for confocal/FIB-SEM overlay. n.s. = no significance; \**P* ≤ 0.05; \*\*\**P* ≤ 0.001; \*\*\*\**P* ≤ 0.0001.

### APOL7C recruitment to phagosomes results in phagosome rupture

To investigate the effect of APOL7C on phagosome membrane integrity we first analyzed the distribution of the membrane damage biomarker galectin 3 in MuTuDC.APOL7C::mCherry cells three hours after phagocytosis. Strikingly, we found that the majority of phagosomes that were positive for APOL7C::mCherry was also positive for galectin 3 (**Figure 3C-D**). This was not the case for APOL7C::mCherry-negative phagosomes, which were rarely positive for galectin 3 (**Figure 3C-D**). To distinguish cause and effect, we performed a gain-of-function experiment in Raw264.7 cells that stably express doxycycline-inducible APOL7C::mCherry (RawKb.APOL7C::mCherry). Raw264.7 cells do not express endogenous APOL7C and so the expression of APOL7C in the RawKb.APOL7C::mCherry cells is strictly under the control of doxycycline. In the absence of doxycycline, we could not detect any damaged (galectin 3-positive) phagosomes in these cells (**Figure 3E-F**). However, upon induction of APOL7C::mCherry there was a significant increase in the number of galectin 3-positive phagosomes (**Figure 3E-F**). Notably, all galectin 3-positive phagosomes were also positive for APOL7C::mCherry (**Figure 3E**). We found similar results when using ALIX as an indirect marker of damaged phagosomes (**Figure 3G-H**). We also assessed whether the appearance of APOL7C::mCherry on phagosomes coincided with the loss of the membrane impermeant dye lucifer yellow (LY) from the lumen of phagosomes. LY was loaded into phagosomes and imaged in live cells 3 hours post phagocytosis. Strikingly, APOL7C::mCherry positive phagosomes lost nearly all luminal LY, likely as a result of phagosome rupture (**Figure 3I-J**). Live-cell confocal imaging of individual phagosomes showed that the appearance of APOL7C::mCherry on phagosomes was followed by the near complete loss of luminal LY within a matter of minutes (**Figure 3K-L**). In addition to the loss of LY from the lumen, we also assessed whether phagosomes lose luminal protons (H^+^) as a result of APOL7C::mCherry recruitment. Notably, phagosomes positive for APOL7C::mCherry had a near neutral pH in stark contrast to APOL7C::mCherry-negative phagosomes, which were acidic, as expected (**Supplementary Figure 4A-C**). To investigate whether in addition to a small membrane impermeant dye such as LY, proteins could also move freely across APOL7C::mCherry-positive phagosome membranes, we generated a cell line that stably expressed cytosolic GFP (cGFP) (henceforth RawKb.cGFP.APOL7C::mCherry cells). In these cells, we readily observed the movement of cGFP from the cytosol to the lumen of APOL7C::mCherry-positive phagosomes but not APOL7C::mCherry negative phagosomes (**Figure 3M-N**). This movement was continuous as we readily observed the continued influx of cGFP in the APOL7C::mCherry positive phagosomes after photobleaching, which was not due to damage from photobleaching as it was not observed with photobleached APOL7C::mCherry-negative phagosomes (**Figure 3O**). We next assessed whether we could detect the loss of the model antigen OVA from APOL7C::mCherry positive phagosomes. To this end, we covalently labeled OVA with the fluorophore AF-633 using a succinimidyl ester derivative if AF-633 and attached the resulting fluorescent OVA to beads using the heterobifunctional crosslinker, cyanamide. After a 4 hour incubation of these beads with RawKb.cGFP.APOL7C::mCherry cells, we readily observed the loss of fluorescent OVA from APOL7C::mCherry positive phagosomes but not APOL7C::mCherry negative phagosomes (**Supplementary Figure 4D-E**). Notably, in the same phagosomes where fluorescent OVA was lost we observed the concomitant influx of cGFP indicative of the non-selective bidirectional movement of proteins across APOL7C::mCherry positive phagosomal membranes (**Supplementary Figure 4D-E**). Next, to formally rule out that APOL7C::mCherry is recruited to already damaged phagosomes, we treated RawKb.APOL7C::mCherry cells with LLOMe to instigate damage of all phagosomes. Even at concentrations of LLOMe that result in the loss of luminal LY from all of the phagosomes in a cell, there was no change in the number of APOL7C::mCherry-positive phagosomes (**Supplementary Figure 4F-G**).

Finally, we examined the ultrastructure of APOL7C::mCherry positive phagosomes by correlative confocal and focused ion beam scanning electron microscopy (FIB-SEM). This revealed discontinuity in the phagosomal membrane that would allow for the escape of the luminal contents of phagosomes to the cytosol, as observed above (**Figure 3P** and **Supplementary Movie 1**). Altogether, the appearance of the membrane damage biomarkers galectin 3 and ALIX, the sudden loss of luminal membrane impermeant dyes, the inability to retain H^+^, and the movement of proteins across the phagosomal membrane, suggest that the recruitment of APOL7C::mCherry to phagosomes results in phagosome rupture.

### APOL7C-dependent phagosome rupture drives XP

We next assessed whether APOL7C-dependent phagosome rupture contributes to XP. We first performed gain-of-function studies. For this, the RawKb.APOL7C::mCherry cell line was used as it allows for the dox-inducible expression of APOL7C::mCherry in the absence of endogenous APOL7C. RawKb cells stably express H-2K^b^ and can therefore present the OVA peptide SIINFEKL to reporter B3Z cells. We ensured that doxycycline itself had no effect on the XP of OVA-beads or on the direct presentation of SIINFEKL peptide by RawKb cells. Indeed, RawKb cells incubated with OVA-beads for 4 hours in the presence or absence of doxycycline showed equivalent capacity to XP (**Figure 4A**). Doxycycline also had no effect on the ability of RawKb cells pulsed with SIINFEKL peptide to elicit B3Z cell activation (**Figure 4B**). However, RawKb.APOL7C::mCherry cells incubated with OVA-beads for 4 hours showed a large increase in XP when treated with doxycycline (**Figure 4C**). Direct presentation of SIINFEKL peptide by RawKb.APOL7C::mCherry cells was unaffected by doxycycline treatment (**Figure 4D**). Importantly, doxycycline did not affect the ability of RawKb.APOL7C::mCherry cells to internalize OVA-beads and did not affect the surface expression of H-2K^b^ (**Supplementary Figure 5A-D**). These data indicate that APOL7C can promote XP in non-cDC1 cells.

**Figure 4.**
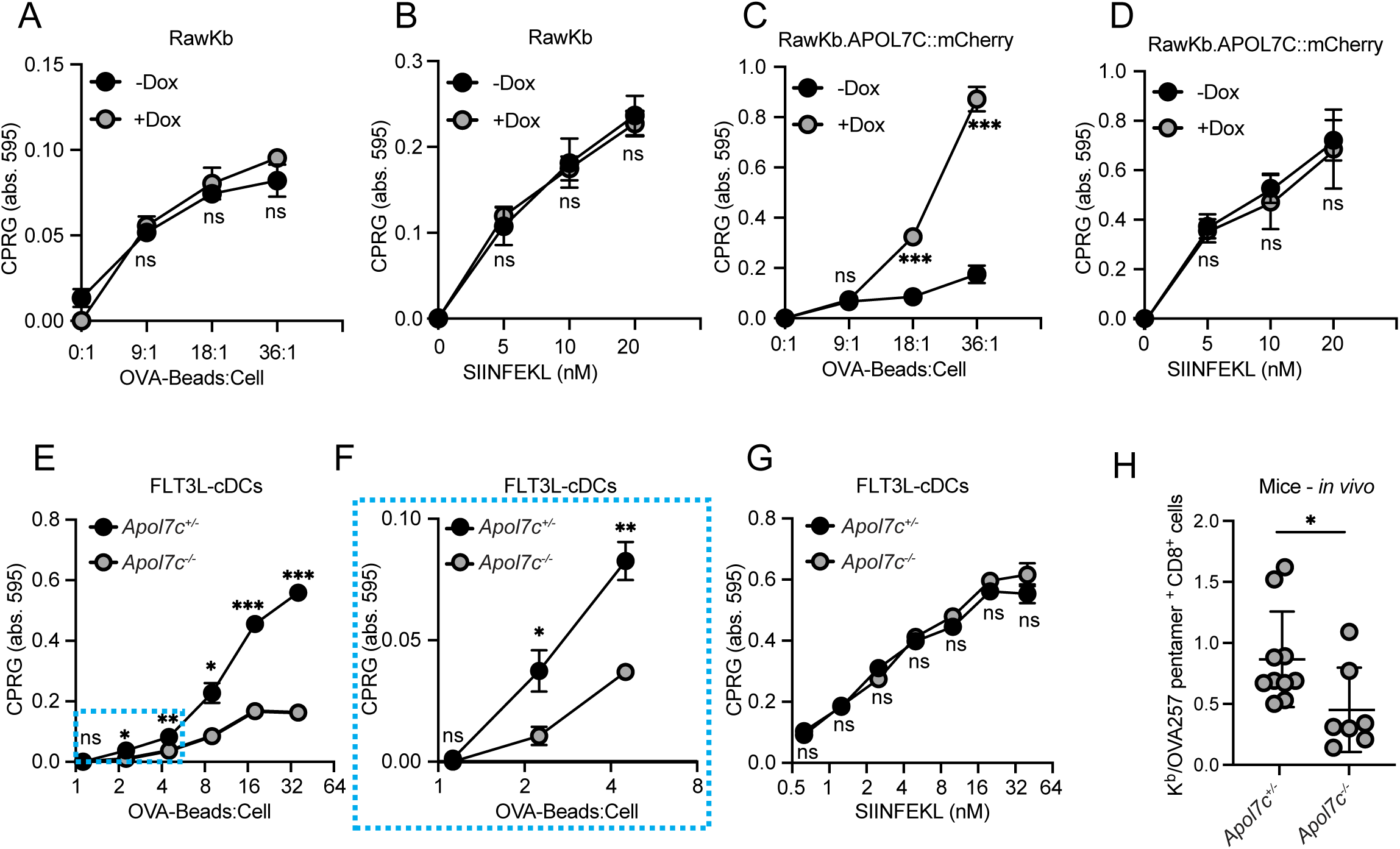
APOL7C expression promotes XP. **(A-D)** RawKb or RawKb.APOL7C::mCherry cells were treated with or without doxycycline (DOX) overnight and then challenged with OVA-beads **(A,C)** or SIINFEKL **(B,D)** for 4hr. The cells were fixed and incubated overnight with B3Z cells. B3Z cells were lysed in a CPRG containing buffer. Plotted as mean ± s.e.m. of experimental triplicates and is representative of 3 independent experiments (*n* = 3). Significance determined using student’s *t* test for individual dilutions. **(E-G)** FLT3L-cDCs derived from littermate *Apol7c^+/-^* and *Apol7c^-/-^* mice were treated with poly(I:C) at 5 μg/ml overnight. After a 4hr incubation with OVA-beads **(E-F)** or SIINFEKL **(G)**, the cells were fixed and incubated overnight with B3Z cells. B3Z cells were lysed in a CPRG containing buffer. Plotted as mean ± s.d. of experimental triplicates and is representative of 3 independent experiments (*n* = 3). Significance determined using student’s *t* test for individual dilutions. **(H)** *Apol7c^+/-^* (n = 10) and *Apol7c^-/-^* (n = 7) littermate mice were immunized i.v. with OVA-beads + 50 μg of poly(I:C). 4-5 days later, the frequency of OVA-specific CD8^+^ T cells in the spleen was determined. Each dot is an individual mouse. Data are represented as mean ± s.d. Data are pooled from three (n = 3) independent experiments. Significance determined using Kolmogorov-Smirnov test. n.s. = no significance; \**P* ≤ 0.05; \*\**P* ≤ 0.01; \*\*\**P* ≤ 0.001.

Next, we performed loss-of-function studies in FLT3L-cDCs derived from wild-type and *Apol7c^-/-^*mice. The *Apol7c^-/-^* mice were generated by removing exons 4 and 5 by CRISPR/Cas9 genome editing (**Supplementary Figure 6A-E**). *Apol7c^-/-^* mice were viable and born at normal Mendelian ratios. Immune profiling of littermate *Apol7c^+/-^* and *Apol7c^-/-^* mice revealed normal myeloid and lymphoid composition, including similar amounts of cDC1s and cDC2s (**Supplementary Figure 7**). However, *Apol7c^-/-^* FLT3L-cDCs had a striking defect in their ability to cross-present OVA-beads (**Figure 4E-F**). Importantly, there was no defect in their ability to directly present SIINFEKL peptide (**Figure 4G**) or in the surface expression of the MHC-I molecule H-2K^b^ (**Supplementary Figure 5E-I**).

Finally, to assess the importance of APOL7C for XP *in vivo*, we immunized *Apol7c^+/-^* and *Apol7c^-/-^* mice with OVA-beds + poly(I:C) and measured OVA-specific CD8^+^ T cell responses by H-2K^b^-OVA-pentamer staining. *Apol7c^+/-^* mice mounted a robust OVA-specific CD8^+^ T cell response that was significantly reduced in *Apol7c^-/-^* littermates (**Figure 4H**). Together, these data indicate that APOL7C can promote the XP of OVA-beads both *in vivo* and *in vitro* by primary FLT3L-cDCs and when ectopically expressed in other phagocytes.

### Apol7C recruitment is independent of SYK but requires NADPH oxidase activity on phagosomes

Phagosome rupture can be modulated by dedicated cDC1 receptors, such as DNGR-1, through a SYK- and NADPH oxidase-dependent pathway^11,22,50,63^. While the OVA beads used here do not harbor DNGR-1 ligands, we wondered whether the recruitment of APOL7C to phagosomes also relied on SYK- and NADPH oxidase-dependent signaling. We stained RawKb.APOL7C::mCherry cells that had phagocytosed OVA beads for phospho-SYK and while we could detect the presence of phospho-SYK on APOL7C::mCherry positive phagosomes (**Figure 5A-B**), inhibition of SYK with the inhibitor R406 had no effect on the recruitment of APOL7C::mCherry (**Figure 5C-D**). We also generated SYK CRISPR knockouts from the RawKb.APOL7C::mCherry cells and observed no difference in the recruitment of APOL7C::mCherry to phagosomes (**Supplementary Figure 8A-C**).

**Figure 5.**
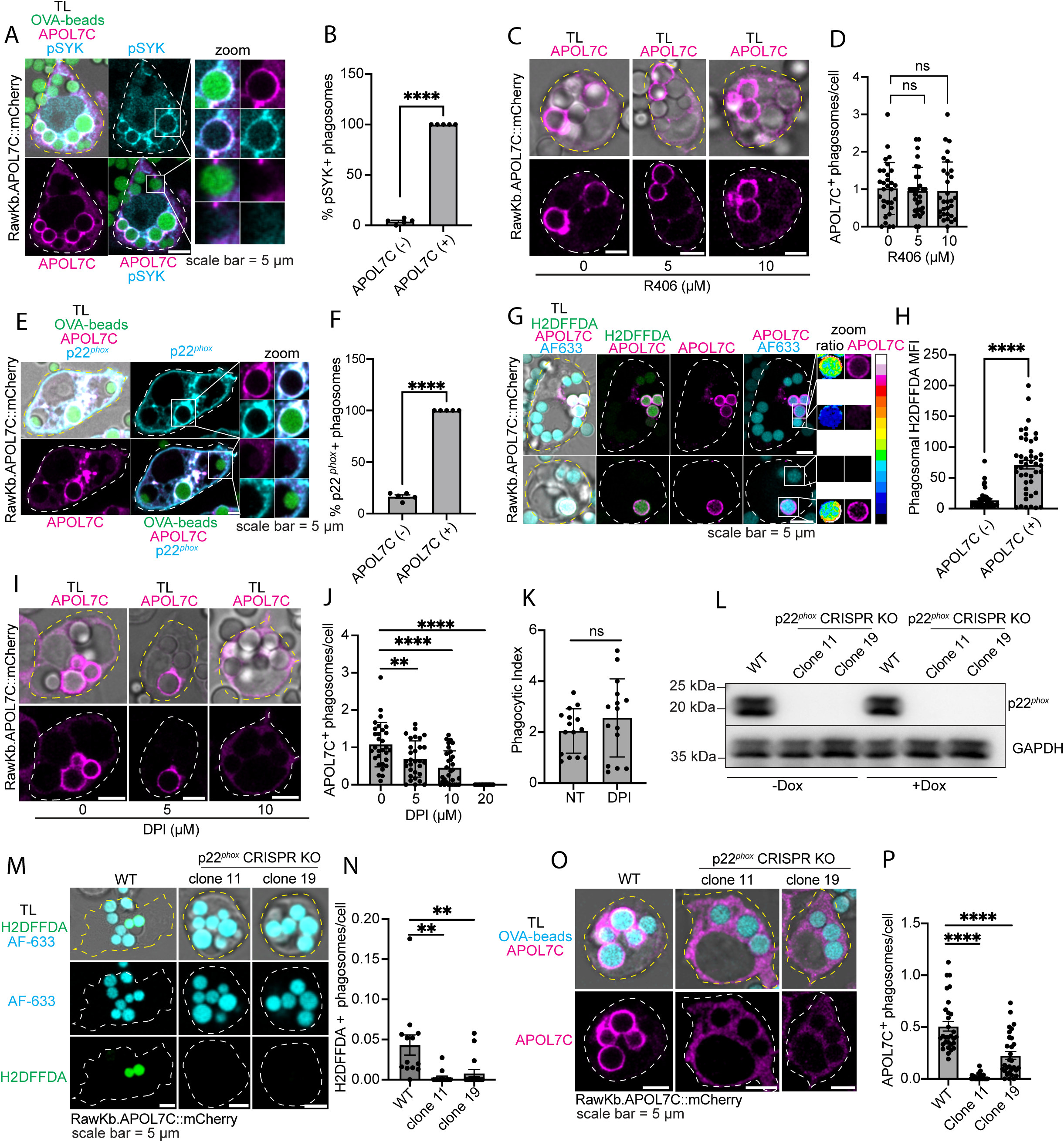
APOL7C recruitment to phagosomes requires NADPH oxidase activity. **(A-B)** RawKb.APOL7C::mCherry cells were treated with doxycycline (DOX) overnight and then challenged with OVA-beads for 4hr. They were then fixed and stained for phosphorylated SYK (pSYK) before imaging by confocal microscopy. Scale bar = 5 μm. The percentage of APOL7C^+^ or APOL7C^-^ phagosomes that were also positive for pSYK are plotted as mean ± s.e.m. Each dot represents a field-of-view containing 5-20 cells (*n* = 5 independent acquisitions). Significance determined using student’s *t* test. **(C-D)** RawKb.APOL7C::mCherry cells were treated with doxycycline (DOX) overnight and then challenged with OVA-beads for 4hr in the presence of R406 at the indicated concentration. Scale bar = 5 μm. The number of APOL7C^+^ phagosomes is plotted as the mean ± s.d. Each dot represents a field-of-view containing 5-20 cells (*n* = 3 independent experiments). Significance determined using student’s *t* test. **(E-F)** RawKb.APOL7C::mCherry cells were treated with doxycycline (DOX) overnight and then challenged with OVA-beads for 4hr. They were then fixed and stained for p22*^phox^* before imaging by confocal microscopy. Scale bar = 5 μm. The percentage of APOL7C^+^ or APOL7C^-^ phagosomes that were also positive for p22*^phox^* are plotted as mean ± s.e.m. Each dot represents a field-of-view containing 5-20 cells (*n* = 5 independent acquisitions). Significance determined using student’s *t* test. **(G-H)** RawKb.APOL7C::mCherry cells were treated with doxycycline (DOX) overnight and then challenged for 4hr with OVA-beads that were dually labeled with the ROS-insensitive dye Alexa Fluor 633 (AF633) and the ROS-sensitive dye H2DFFDA. The cells were then imaged live by confocal microscopy. Pseudocolor and color map indicate degree of bead oxidation. Scale bar = 5 μm. The MFI of H2DFFDA for APOL7C^+^ or APOL7C^-^ phagosomes is plotted as mean ± s.e.m. Each dot represents a single phagosome. Data pooled from 3 independent experiments (*n* = 3). Significance determined using student’s *t* test. **(I-J)** RawKb.APOL7C::mCherry cells were treated with doxycycline (DOX) overnight and then challenged with OVA-beads for 4hr in the presence of DPI at the indicated concentration. Scale bar = 5 μm. The number of APOL7C^+^ phagosomes is plotted as the mean ± s.d. Each dot represents a field-of-view containing 5-20 cells (*n* = 3 independent experiments). Significance determined using student’s *t* test. **(K)** The number of internalized OVA-beads were counted from the micrographs shown in **(I)**. Plotted as mean ± s.d. Each dot represents a field-of-view containing 5-20 cells (*n* = 3 independent experiments). Significance was determined using a student’s *t* test. **(L)** Western blot showing p22*^phox^* levels in wild-type and p22*^phox^* KO RawKb.APOL7C::mCherry cells. **(M-N)** RawKb.APOL7C::mCherry cells were challenged for 1hr with OVA-beads that were dually labeled with AF633 and H2DFFDA. The cells were then imaged live by confocal microscopy. Scale bar = 5 μm. The number of H2DFFDA^+^ phagosomes is plotted as the mean ± s.e.m. Each dot represents a field-of-view containing 5-20 cells (*n* = 3 independent experiments). Significance determined using student’s *t* test. **(O-P)** The indicated cells were treated with DOX overnight and challenged with OVA-beads for 4hr. Scale bar = 5 μm. The number of APOL7C^+^ phagosomes is plotted as the mean ± s.e.m. Each dot represents a field-of-view containing 5-20 cells (*n* = 3 independent experiments). Significance determined using student’s *t* test. n.s. = no significance; \*\**P* ≤ 0.01; \*\*\*\**P* ≤ 0.0001.

Next we checked whether the NADPH oxidase was present on APOL7C::mCherry positive phagosomes. Staining for the p22*^phox^* subunit of the oxidase revealed that it was also present on APOL7C::mCherry positive phagosomes (**Figure 5E-F**). We could also detect oxidase activity in phagosomes by covalently attaching the reactive oxygen species (ROS)-sensitive dye H2DFFDA along with a ROS-insensitive fluorophore Alexa Fluor 633 (AF633) directly to the beads. Oxidation of H2DFFDA in the phagosomes was sensitive to the NADPH oxidase inhibitor diphenylene iodonium (DPI) (**Supplementary Figure 8D-E**). As previously reported^64^, we observed heterogeneity in the ability of individual phagosomes to generate superoxide (**Figure 5G**). Interestingly, we found a very strong correlation between oxidase activity and the recruitment of APOL7C::mCherry to phagosomes suggesting a potential role for the NADPH oxidase in the recruitment of APOL7C::mCherry (**Figure 5G-H**). DPI treatment of the RawKb.APOL7C::mCherry cells revealed a dose-dependent reduction in APOL7C::mCherry recruitment, without any effect on OVA bead uptake (**Figure 5I-K**). We also generated p22*^phox^* CRISPR knockouts from the RawKb.APOL7CmCherry line which were incapable of producing phagosomal ROS (**Figure 5L-N**). The knockout cell lines also showed a significant decrease in the number of APOL7C::mCherry positive phagosomes (**Figure 5O-P**). Altogether our data indicate that APOL7C recruitment to OVA bead-containing phagosomes is independent of SYK but does require the activity of the NADPH oxidase.

### NADPH oxidase activity is required for APOL7C-dependent XP

As discussed earlier, expression of APOL7C::mCherry in cells that normally lack APOL7C results in enhanced XP. We next investigated whether NADPH oxidase activity, which appears to be a pre-requisite for APOL7C recruitment to phagosomes, is necessary for APOL7C-dependent XP. We first tested the effect of acute inhibition of oxidase activity with DPI. Treatment of RawKb.APOL7C::mCherry cells with DPI completely inhibited the enhanced XP observed upon APOL7C::mCherry expression without having any effect on the direct presentation of SIINFEKL peptide, OVA bead uptake, or on the surface expression of H-2K^b^ (**Supplementary Figure 9A-C**). Similarly, unlike the parental RawKb.APOL7C::mCherry cells (**Figure 4K-L** and **Supplementary Figure 9D-F**), both of the p22*^phox^* knockout clones generated above failed to show enhanced XP upon doxycycline induction of APOL7C::mCherry expression despite robust upregulation of APOL7C::mCherry (**Supplementary Figure 9G and 9J**). This was not due to changes to their ability to directly present SIINFEKL peptide (**Supplementary Figure 9H and 9K**), or to changes in the surface expression of H-2K^b^ (**Supplementary Figure 9I and 9L**). Together these findings indicate that APOL7C-dependent XP is dependent upon NADPH oxidase activity.

### APOL7C insertion into phagosomes is a voltage-dependent process that relies on NADPH oxidase-driven depolarization of the phagosome membrane

The rupture of endocytic organelles, including phagosomes, in cDC1s has been proposed to occur as a result of peroxidation of their limiting lipid bilayer by NADPH oxidase-derived ROS^27,65^. It was conceivable, however, that lipid peroxidation of the phagosomal membrane may result in the insertion of APOL7C. For this, we employed the radical-trapping antioxidant liproxstatin-1 (LPX) which potently inhibits the accumulation of lipid peroxides in cellular membranes^66,67^. LPX treatment had no effect on the uptake of OVA beads, nor did it affect the oxidation of H2DFFDA on the OVA beads (**Figure 6A-C**). It did however potently inhibit the oxidation of BODIPY 581/591 C11, a redox-sensitive lipid oxidation probe (**Figure 6D-E**). This contrasts with DPI which inhibits the enzymatic activity of the NADPH oxidase and therefore blocks oxidation of both H2DFFDA and BODIPY 581/591 C11 (**Figure 6A-E**). Notably, unlike DPI, LPX treatment had no effect on either the recruitment of APOL7C::mCherry to phagosomes or on APOL7C-dependent XP (**Figure 6F-J**). This indicates that lipid peroxidation is unlikely to be the signal for APOL7C insertion into phagosome membranes.

**Figure 6.**
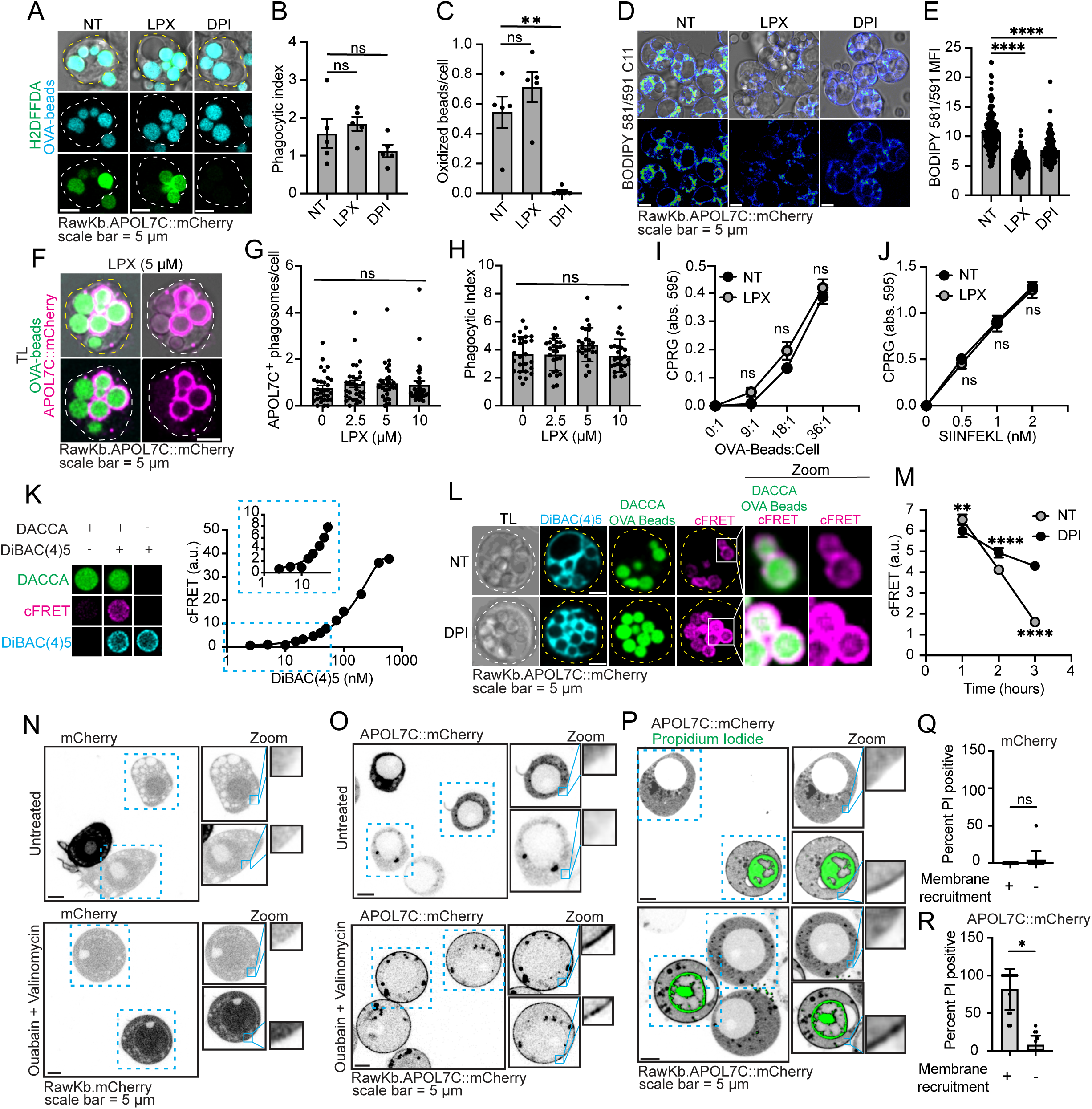
APOL7C recruitment to phagosomal membranes is not dependent of oxidase-driven lipid peroxidation but is voltage-dependent. **(A)** RawKb.APOL7C::mCherry cells were treated with liproxstatin 1 (LPX) or DPI and challenged with AF633- and H2DFFDA-labeled OVA-beads. Images acquired by confocal microscopy. Scale bar = 5 μm. **(B)** The number of OVA-beads per cell is plotted as the mean ± s.e.m. Each dot represents a field-of-view containing 5-20 cells (*n* = 3 independent experiments). Significance determined using Kolmogorov-Smirnov test. **(C)** The number of oxidised (H2DFFDA^+^) beads per cell is plotted as the mean ± s.e.m. Each dot represents a field-of-view containing 5-20 cells (*n* = 3 independent experiments). Significance determined using Kolmogorov-Smirnov test. **(D-E)** RawKb.APOL7C::mCherry cells were preloaded with the lipid peroxidation probe BODIPY 581/591 C11 before a 4hr challenge with OVA-beads in the presence of the indicated drugs. Images were acquired by live cell confocal microscopy with excitation set to 488 nm and emission to 510 nm. Scale bar = 5 μm. Plotted is the mean ± s.d. of the MFI (exc. 488/ems. 510) of individual cells from 3 independent experiments (*n* = 3). Significance determined using Kolmogorov-Smirnov test. **(F-G)** RawKb.APOL7C::mCherry cells were treated with doxycycline (DOX) overnight and then challenged with OVA-beads for 4hr in the presence of LPX at the indicated concentration. Scale bar = 5 μm. The number of APOL7C^+^ phagosomes is plotted as the mean ± s.e.m. Each dot represents a field-of-view containing 5-20 cells (*n* = 3 independent experiments). Significance determined using student’s *t* test. **(H)** The number of internalized OVA-beads were counted from the micrographs shown in **(F)**. Plotted as mean ± s.e.m. Each dot represents a field-of-view containing 5-20 cells (*n* = 3 independent experiments). Significance was determined using a student’s *t* test. **(I-J)** RawKb.APOL7C::mCherry cells were treated with or without doxycycline (DOX) overnight and then challenged with OVA-beads **(I)** or SIINFEKL **(J)** for 4hr. The cells were fixed and incubated overnight with B3Z cells. B3Z cells were lysed in a CPRG containing buffer. Plotted as mean ± s.e.m. of experimental triplicates and is representative of 4 independent experiments (*n* = 4). Significance determined using student’s *t* test for individual dilutions. **(K)** OVA-beads labelled with DACCA were bathed in buffers containing increasing concentrations of DiBAC(4)5. Plotted are the corrected FRET (cFRET) values ± s.e.m. for each concentration. Number of beads analyzed = 249-1267. **(L-M)** RawKb.APOL7C::mCherry cells were challenged with DACCA-labeled OVA-beads for the indicated time and at the time of confocal imaging, DiBAC(4)5 was added to the cells. cFRET was assessed for individual phagosomes. Plotted is the mean ± s.e.m. of the corrected cFRET signal from individual phagosomes (*n* = ≥ 500 phagosomes per condition). **(N-R)** RawKb.mCherry **(N, Q)** or RawKb.APOL7C::mCherry **(O, P, R)** were treated with ouabain for 3hr followed by a treatment with valinomycin in K^+^-rich, Na^+^-free medium containing propidium iodide (PI) **(P,Q,R)** or not **(N,O)** for 30min. Images were acquired by live cell confocal microscopy. Scale bar = 5 μm. The percentage of PI^+^ cells is plotted as the mean ± s.d. Each dot represents a field-of-view containing 5-20 cells (*n* = 3 independent experiments). Significance determined using student’s *t* test. n.s. = no significance; \**P* ≤ 0.05; \*\**P* ≤ 0.01; \*\*\*\**P* ≤ 0.0001.

We next investigated other consequences of NADPH oxidase activity that may mark phagosomes for APOL7C insertion. In addition to generating ROS, the NADPH oxidase has electrogenic activity^68–70^. This is a consequence of the transfer of electrons from a cytosolic electron donor, NADPH, to an electron acceptor, O_2,_ generating O_2_^-^ that is released into the phagosomal lumen. The catalytic subunit of the NADPH oxidase enzyme complex spans the membrane and transfers electrons across the bilayer thereby generating a transmembrane voltage. At the plasma membrane (PM) of neutrophils, for example, NADPH oxidase activity can result in depolarization of the PM from roughly -60 mV to +58 ± 6 mV^68^. Interestingly, several members of the human APOL family insert into lipid bilayers in a voltage-dependent manner^71^. Notably, APOL3, APOL4 and APOL5 have been shown to insert into lipid bilayers in a voltage-dependent manner, and require a positive voltage on the side of insertion^71^. This contrasts with APOL1 and APOL2, which insert into bilayers in a pH-dependent manner. The voltage-gated insertion of APOL3-5 has been mapped to three key residues found in a putative transmembrane domain called the membrane insertion domain (MID)^71,72^ (**Supplementary Figure 6B** and **Figure 7A**). These residues are also present in APOL7C and correspond to A178, A186 and G190 of the APOL7C MID (**Figure 7A**). It is therefore conceivable that NADPH oxidase-driven depolarization of the phagosome membrane may be the trigger for the insertion of APOL7C into phagosome membranes. To investigate this possibility, we first measured the membrane potential of phagosomes in untreated cells and in cells treated with DPI. Phagosomal membrane potential measurements were performed using a previously established FRET-based approach^73^. OVA beads were covalently labeled with the succinimidyl ester of the FRET donor 7-diethlyaminocoumarin-3-carboxylic acid (DACCA). The potential-sensitive, lipid-soluble probe *bis*-(1,3-dibutylbarbituric acid)pentamethine oxonol [DiBAC_4_(5)], which partitions across membranes in accordance with their transmembrane potential was used as the FRET acceptor. As FRET relies on the close apposition of the donor:acceptor pair (< 10 nm), the physical attachment of the donor fluorophore to the phagocytic target restricts the FRET signal to the phagosome. Bathing DACCA-OVA-beads in increasing concentrations of DiBAC_4_(5) revealed that the corrected FRET (cFRET) signal was restricted to the OVA-bead surface and that sensitive cFRET measurements could be attained over a broad range of DiBAC_4_(5) concentrations (**Figure 6K**). Next we fed DACCA-OVA-beads to RawKb.APOL7C::mCherry cells for 1 hour, then thoroughly washed away external beads, and bathed the cells in 250 nM DiBAC_4_(5). We found that the DiBAC_4_(5) was, as expected, present throughout the cell but the cFRET signal was restricted to the phagosome (**Figure 6L**). As the NADPH oxidase transfers electrons from the cytosol to the lumen of the phagosome, oxidase activity would result in the exclusion of DiBAC_4_(5), which has a negative charge, from the phagosome and therefore a reduced cFRET signal. Interestingly, we found a progressive depolarization (reduced phagosomal cFRET) of the phagosome membrane over the first 3 hours post-phagocytosis (**Figure 6M**). This depolarization was DPI-sensitive, indicating that it was likely driven by NADPH oxidase activity (**Figure 6M**). It is therefore possible that NADPH oxidase-driven depolarization of the phagosomal membrane can instigate insertion of APOL7C.

**Figure 7.**
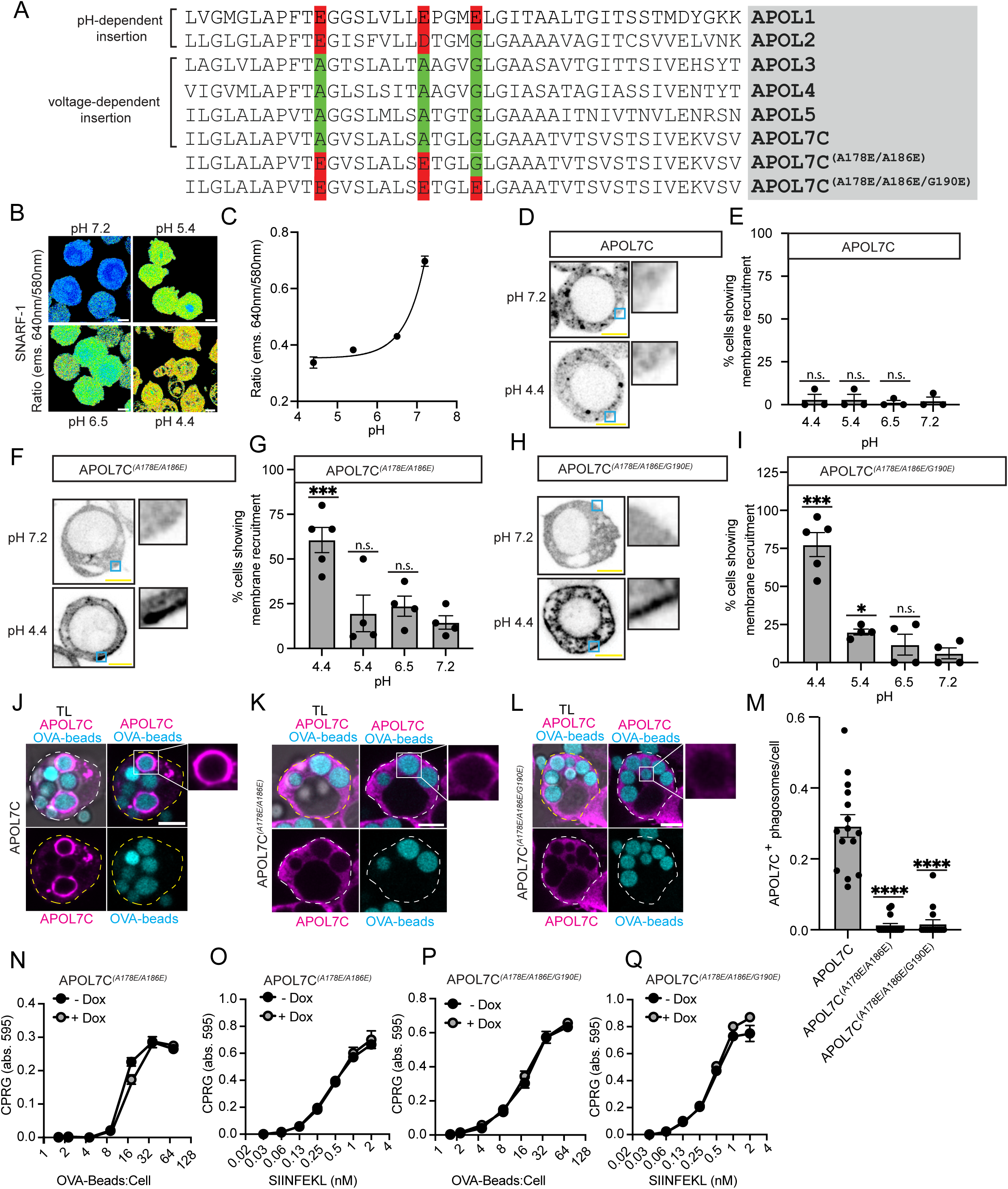
Voltage-dependence of APOL7C-dependent phagosomal recruitment maps to two key residues in the membrane insertion domain. **(A)** Alignment of APOL7C (residues 168-210) with APOL1-6. Also shown is alignment with the APOL7C^(A178E/A186E)^ and APOL7C^(A178E/A186E/G190E)^ mutants. Scale bar = 5 μm. **(B-C)** Ratiometric imaging of SNARF-1 upon clamping with at the indicated pH. pH was clamped by bathing the cells in a K^+^-rich medium, containing 10 μg/ml nigericin, at the indicated pH. Plotted is mean ± s.d. of the ems. 640nm/580nm ratio (*n* = 30 cells per condition). **(D-I)** RawKb.APOL7C::mCherry, RawKb.APOL7C^(A178E/A186E)^::mCherry, or RawKb. APOL7C^(A178E/A186E/G190E)^::mCherry cells were clamped at the indicated pH and imaged by live cell confocal microscopy. Scale bars = 5 μm. The number of cells showing redistribution of APOL7C::mCherry to the plasma membrane is plotted as the mean ± s.e.m. Each dot represents a field-of-view containing 5-20 cells (*n* = 3-5 independent experiments). Significance was determined using a student’s t test. **(J-M)** RawKb.APOL7C::mCherry, RawKb.APOL7C^(A178E/A186E)^::mCherry, or RawKb. APOL7C^(A178E/A186E/G190E)^::mCherry cells were incubated with doxycycline overnight and then challenged with OVA-beads for 4hr before confocal imaging. Plotted is the number of APOL7C^+^ phagosomes per cell. Each dot represents a field-of-view containing 5-20 cells (*n* = 3 independent experiments). Significance was determined using a student’s t test. **(N-Q)** RawKb.APOL7C::mCherry, RawKb.APOL7C^(A178E/A186E)^::mCherry, or RawKb. APOL7C^(A178E/A186E/G190E)^::mCherry cells were treated with or without doxycycline (DOX) overnight and then challenged with OVA-beads or SIINFEKL for 4hr. The cells were fixed and incubated overnight with B3Z cells. B3Z cells were lysed in a CPRG containing buffer. Plotted as mean ± s.e.m. of experimental triplicates and is representative of 8 independent experiments (*n* = 4). Significance determined using student’s *t* test for individual dilutions. n.s. = no significance; \*\*\**P* ≤ 0.001; \*\*\*\**P* ≤ 0.0001.

To next test the voltage-dependence of APOL7C insertion into lipid bilayers we employed both cellular- and giant unilamellar vesicle (GUV)-based approaches. First, as we could not manipulate phagosome membrane potential without affecting other cellular membranes, we attempted to clamp the plasma membrane (PM) potential of RawKb.mCherry and RawKb.APOL7C::mCherry cells such that the cytosolic side became positive by imposing an inward K^+^ gradient in the presence of a K^+^-selective ionophore. We reasoned that this would expose APOL7C::mCherry to positive voltages on the cytosolic side and therefore drive its insertion into the PM. To achieve this, we loaded the cytosol of the cells with Na^+^ and depleted them of K^+^ by inhibiting Na^+^/K^+^- ATPase activity with ouabain for three hours. Then we transferred the cells to a K^+^-rich, Na^+^-free medium in the presence of the K^+^-selective ionophore valinomycin. This imposes an inward K^+^ gradient and is predicted to clamp the plasma membrane potential (χπ_pm_) at (RT/zF)ln([K^+^*_i_*]/[K^+^*_o_*]) where [K^+^*_i_*] and [K^+^*_o_*] are the cytosolic and extracellular concentrations of K^+^, R, T and F are the gas constant, temperature in degrees kelvin, and Faraday’s constant, respectively, and *z* is the charge of K^+^. This treatment had no effect on the distribution of mCherry but, remarkably, resulted in the near immediate insertion of APOL7C::mCherry into the plasma membrane of the RawKb.APOL7C::mCherry cells (**Figure 6N, 6O** and **6Q**). Moreover, insertion of APOL7C::mCherry into the PM resulted in its permeabilization to propidium iodide (PI) (**Figure 6P** and **6R**).

We next employed a GUV-based approach to investigate the voltage-gated insertion of APOL7C in lipid bilayers. To this end, we generated GUVs containing the monovalent cation-selective ionophore gramicidin^74,75^. By generating these gramicidin-containing GUVs in the presence of a K^+^-rich medium and subsequently varying the K^+^ concentration (while keeping osmolarity constant with tetraethylammonium chloride) in the medium the GUVs are bathed in, the membrane potential across the GUV membrane can be clamped at discrete values described by the Nernst equation. Using the potential-sensitive dye 3,3’-dihexyloxacarbocyanine iodide [DiOC_6_(3)], we first confirmed that varying the K^+^ concentration reliably altered the potential across the GUV membrane (**Supplementary Figure 10A-B**). Next, we investigated whether exposing recombinant APOL7C::GFP to positive voltages on the *cis* (outer) side of the GUV would facilitate its insertion into the GUV membrane. When the potential of the GUV membrane was 0 mV, APOL7C::GFP did not insert into the GUV membrane. However, when the potential of the GUV membrane was 71 mV (outside positive), APOL7C::GFP inserted into the GUV membrane and insertion was accompanied by vesiculation and tubulation of the GUV (**Supplementary Figure 10C-E**).

To further investigate the voltage-dependence of APOL7C insertion into phagosome membranes, we next assessed whether mutation of the 3 key residues that dictate voltage or pH-dependent insertion of APOLs would impact APOL7C insertion into phagosome membranes. As discussed above, APOLs insert into bilayers in either a pH- or voltage-dependent manner. This is largely determined by three residues in the MID that align with E201, E209 and E213 of APOL1^71,72^ (**Figure 7A**). The pH-dependent APOLs, APOL1 and APOL2, have conserved negatively charged amino acids at these positions, whereas all other APOLs, including APOL7C, have neutral amino acids at these residues. Importantly, negatively charged amino acids at these residues not only dictate pH-dependent insertion, but render APOLs voltage-insensitive^71,72^. We therefore reasoned that converting the neutral amino acid residues at positions A178, A186 and G190 of APOL7C (corresponding to E201, E209 and E213 in APOL1) to negatively charged glutamic acid residues would block the voltage-dependent insertion of APOL7C into phagosome membranes. Indeed, we found that APOL7C^(A178E/A186E)^::mCherry and APOL7C^(A178E/A186E/G190E)^::mCherry mutants were not recruited to phagosome membranes (**Figure 7J-M**) and had no impact on XP (**Figure 7N-Q**). This was not due to their inability to insert into membranes, as both APOL7C^(A178E/A186E)^::mCherry and APOL7C^(A178E/A186E/G190E)^::mCherry mutants readily inserted into cellular membranes when the cytosolic pH was clamped at acidic pH; whereas, wild-type APOL7C::mCherry did not (**Figure 7B-I**). Therefore, converting APOL7C to a voltage-insensitive APOL, akin to APOL1 and APOL2, completely blocked both its recruitment to phagosomes and its ability to drive XP.

Altogether our data indicate that lipid peroxidation is unlikely to affect the insertion of APOL7C into the phagosome membrane. Instead, APOL7C insertion into membranes appears to be voltage-dependent and insertion into phagosomes is likely driven by NADPH oxidase-induced depolarization of the phagosome membrane.

## Discussion

XP is important for the generation of many virus- and tumour-specific CTL responses^1–8^. Despite its obvious biological importance, the mechanism(s) by which XP is achieved remain unclear^10,19,63,76–79^. Two recent observations from the XP literature informed the present study. First, while XP has been observed in a variety of APCs *in vitro*, only cDC1s appear to serve a non-redundant role in cross-priming to many types of viruses and tumours *in vivo*^1,9,10,80–83^. Second, the XP of cell-associated antigens by cDC1s often involves the delivery of antigens to the cytosol^11,20,21,23,28,34,50,63^. In line with this, several groups have recently shown that, in cDC1s, endocytic organelles, including phagosomes, appear to be “damage prone” and to display signs of breached or ruptured membranes^11,23,27,28,65^. This is believed to facilitate the delivery of endocytosed or phagocytosed antigens to the cytosol where they gain access to the endogenous pathway of MHC class I processing and presentation^63,76,78^. Given the specialized role of cDC1s in XP *in vivo* and the observation that endocytic organelles in cDC1s often display signs of compromised membrane integrity, we hypothesized that cDC1s harbor adaptations that facilitate the delivery of phagocytosed antigens to the cytosol where they can be processed for XP.

Our data indicate that cDC1s express the cytosolic, pore-forming protein APOL7C. APOL7C is recruited to phagosomes in a regulated manner that requires the NADPH oxidase-dependent depolarization of the phagosomal membrane. Recruitment of APOL7C to phagosomes instigates the formation of large, nonselective pores or breaches in the phagosomal membrane. This in turn allows for the release of phagosomal contents to the cytosol for XP. The unique expression pattern of APOL7C and its role in the delivery of phagocytosed material to the cytosol supports our hypothesis and supports other reports^23^ indicating the existence of dedicated cDC-specific pore-forming proteins that can facilitate XP.

Two main pathways have been described for the escape of phagosomal contents from the phagosomes of APCs during XP – the “transporter” hypothesis and the “indigestion” or phagosomal rupture hypothesis^18,78^. Our findings support the “indigestion” hypothesis. Interestingly, we show that cDC1s not only possess dedicated pore-forming proteins that are recruited to phagosomes to elicit phagosomal rupture, but that the process is tightly regulated. We propose that APOL7C-dependent phagosomal rupture is regulated at least at three different levels. First, we found that the TLR3 agonist poly(I:C) significantly increased the expression of APOL7C. This implies a priming effect of innate immune stimuli and is consistent with the concept of cDC activation whereby cDCs undergo gene transcription and translation changes in response to stimulation to increase their antigen processing and presentation capacity^24–26,36,40^. Second, we found that not all phagosomes recruit APOL7C. Instead, APOL7C is selectively recruited to phagosomes on which the NADPH oxidase is active. Although the heterogeneity of NADPH oxidase activity at the level of individual phagosomes has been previously documented^64^, the basis of this heterogeneity is poorly understood. We speculate that receptors present in the phagosome survey its contents and trigger oxidase activity. This notion is supported by the recent finding that DNGR-1, a cDC1-specific receptor that signals for phagosome rupture, can drive NADPH oxidase activity in phagosomes where DNGR-1 ligands are present^11,22^. We cannot, however, rule out that other, non-cargo dependent pathways contribute to the observed heterogeneity in phagosomal oxidase activity and the subsequent recruitment of APOL7C. Finally, cellular membrane repair pathways are also likely to regulate APOL7C-dependent phagosome rupture. We observed the preferential recruitment of the ESCRT-III machinery and galectin 3 to APOL7C-positive phagosomes. Both have been described to repair damaged endocytic organelles including phagosomes^46–49,51,54^. Notably, genetic ablation of components of the ESCRT-III machinery has been shown to increase XP in cDC1s^28^. Altogether our findings, and others, support the notion that APOL7C-dependent phagosomal rupture is a carefully regulated process.

While we present clear evidence for APOL7C-dependent phagosome rupture, the nature of the damage inflicted to phagosome membranes remains unclear. At least three possibilities exist. First, several members of the human apolipoprotein L family (APOLs) have been shown to serve as channels for cations, namely Na^+^ and K^+^ ^55,71,72^. It is possible that ion fluxes across the phagosomal membrane may impose osmotic stress leading to rupture. The extent and nature of the latter is difficult to deduce, as precise determinations of the luminal ion concentrations of phagosomes are not available. Second, the human apolipoprotein L (APOL3) was recently shown to possess detergent-like properties^58^. APOL3 is capable of dissolving anionic membranes, such as the membranes of cytosol-invading bacteria. Notably, we also observed the rapid dissolution of GUV membranes in the presence of recombinant APOL7C. However, this does not explain why other cellular membranes, such as the plasma membrane and endosomes, which are also abundant in negatively charged lipids such as phosphatidylserine and phosphoinositide species are not similarly targeted. Clearly, other regulatory mechanisms exist. Finally, several APOLs have been shown to oligomerize via a c-terminal leucine zipper. Oligomerization of APOLs appears to be critical for their ion-conducting properties. The nature of oligomeric APOL complexes is currently unclear but, akin to other pore-forming proteins^84,85^, they may possess membrane-disrupting properties.

Altogether, our data identify dedicated pore-forming apolipoproteins that operate in the endocytic pathway of cDCs. We find that one of these apolipoproteins, APOL7C, supports XP by mediating the delivery of exogenous antigen to the cytosol. While we show that the expression of APOL7C is inducible and that its recruitment to phagosomes is dependent on the NADPH oxidase, future studies on what signals drive oxidase activity and therefore “mark” individual phagosomes for APOL7C recruitment and phagosome rupture are of considerable interest and may inform immunotherapeutic strategies for priming CD8^+^ T cells in the context of cancer treatment and vaccines.

## Figure Legends

**Supplementary figure 1.**
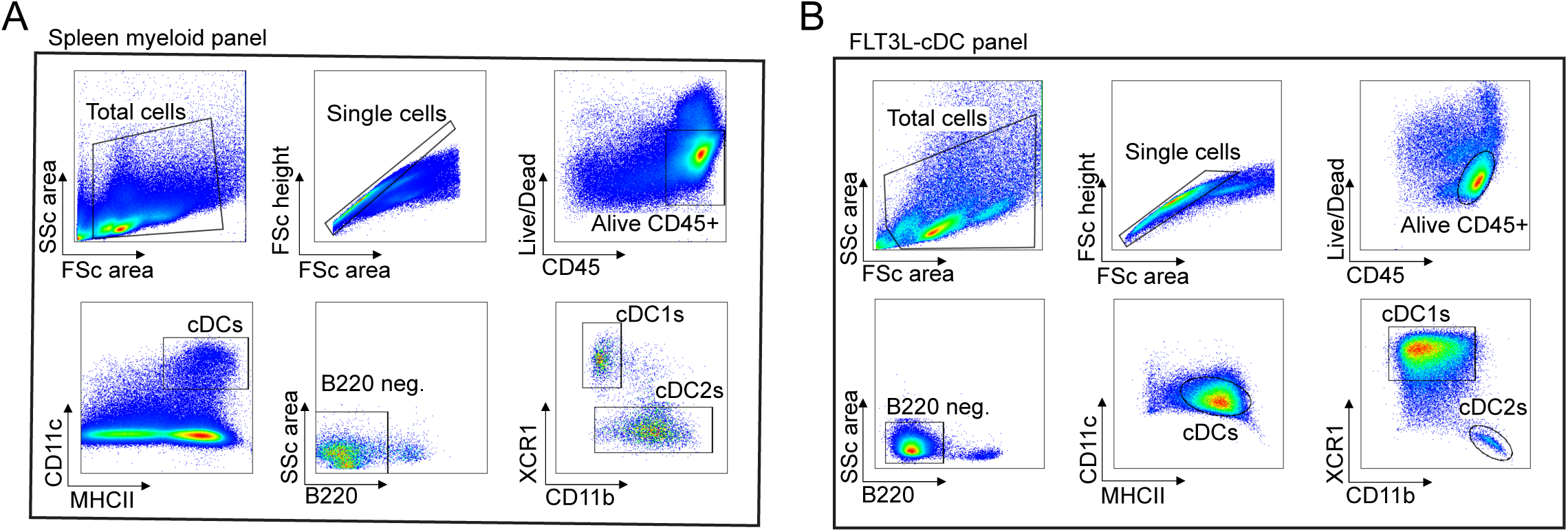
Gating strategy for cDCs. Related to Figure 1 and Figure 4. **(A-C)** Single cell suspensions from the spleen or FLT3L-cDCs were stained and their composition assessed by flow cytometry. cDC1s are considered to be CD45^+^, B220^-^, CD11c^hi^, MHC-II^hi^, CD11b^-^, and XCR1^+^. cDC2s are considered to be CD45^+^, B220^-^, CD11c^hi^, MHC-II^hi^, CD11b^+^, and XCR1^-^. Shown are representative plots from 2-4 similar experiments (*n* = 2-4).

**Supplementary figure 2.**
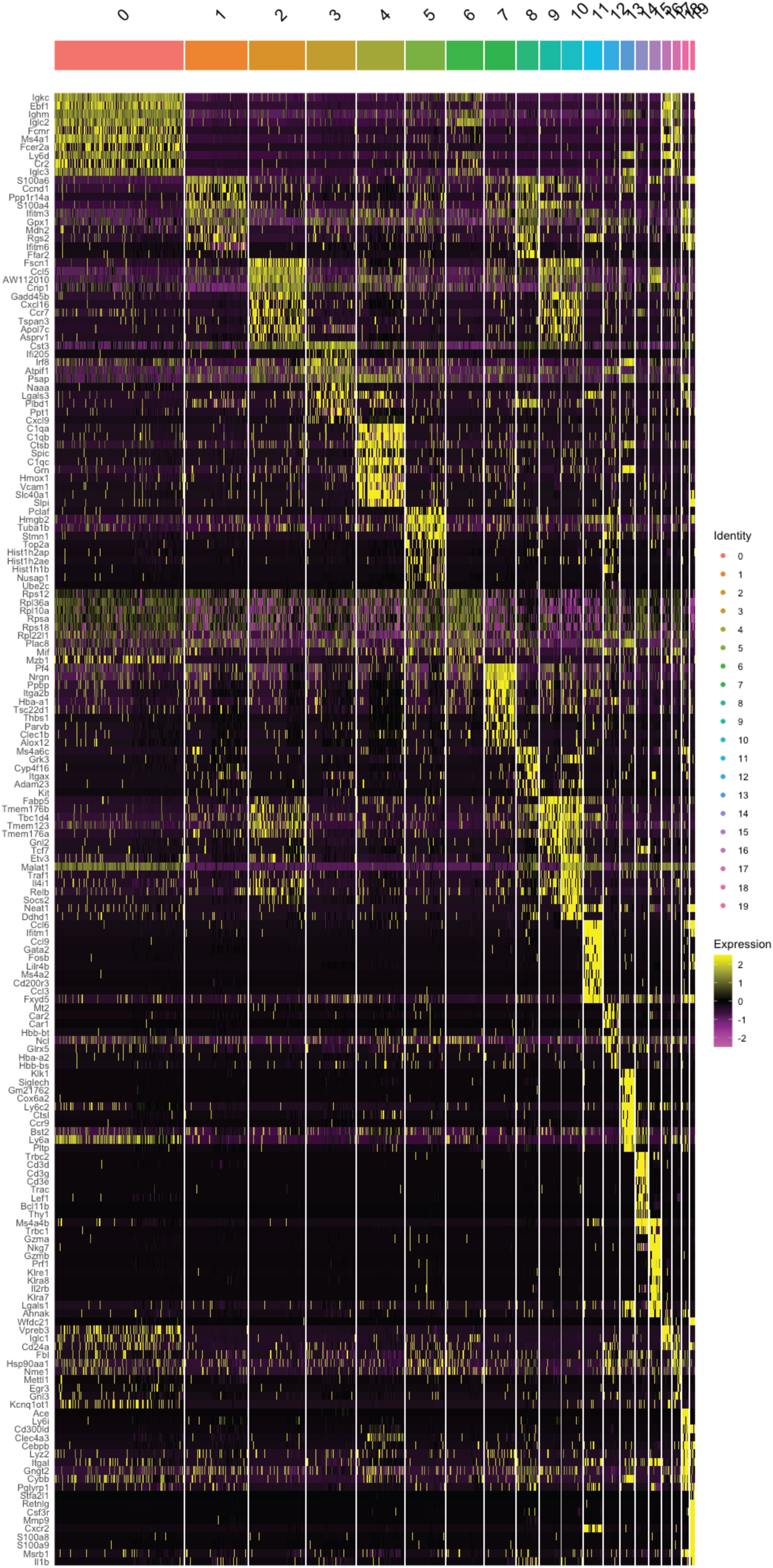
Heatmap of top 10 differentially expressed genes (DEGs) across all clusters. Related to Figure 2. Shown is a heatmap of the top 10 DEGs across all groups from scRNAseq. Colors indicate average expression of each gene.

**Supplementary figure 3.**
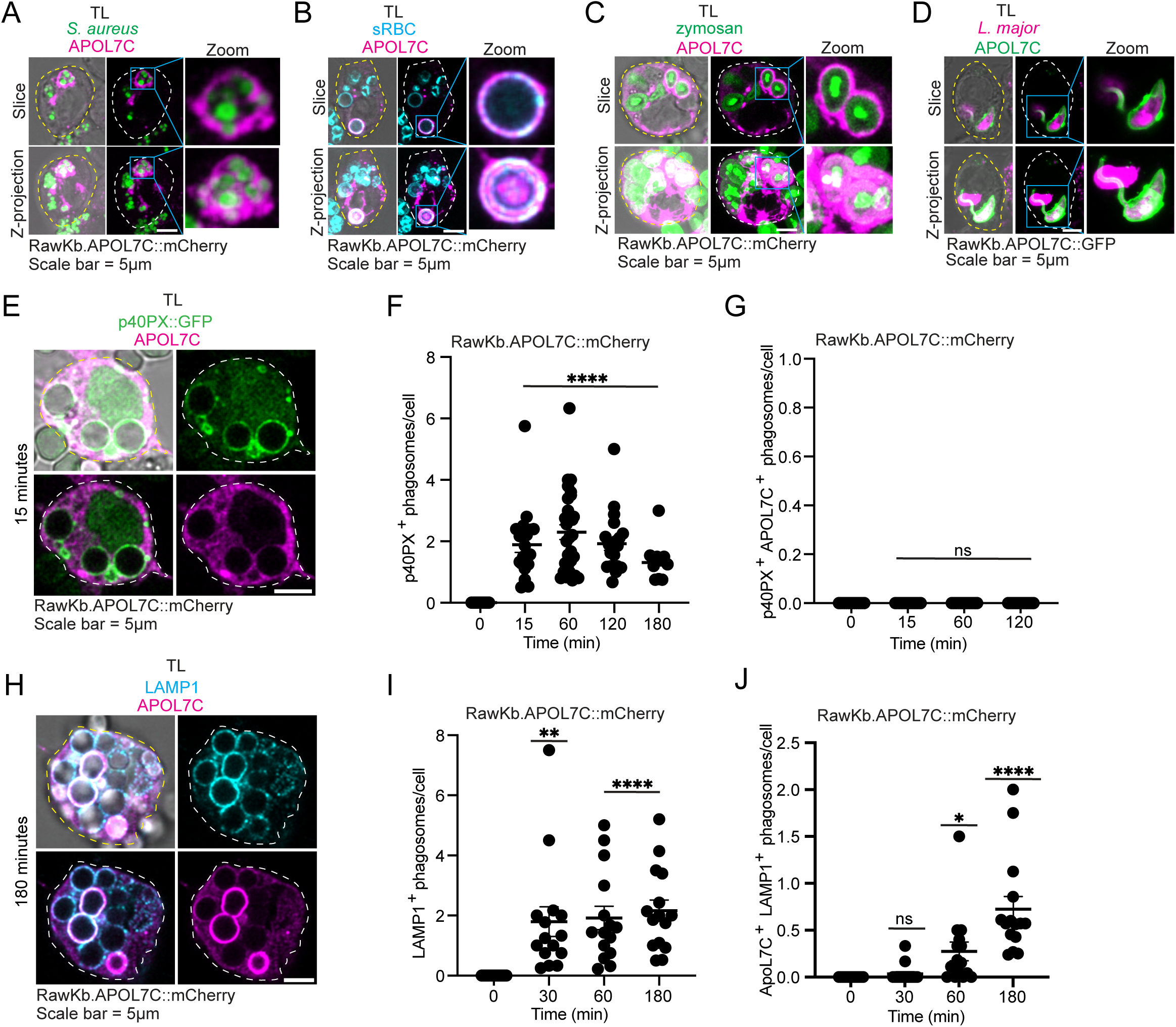
APOL7C::mCherry is recruited to LAMP1^+^ phagosomes harboring different types of cargo. Related to Figure 3. **(A-D)** RawKb.APOL7C::mCherry **(A-C)** or RawKb.APOL7C::GFP **(D)** cells were challenged with *Staphylococcus aureus* **(A)**, IgG-opsonized sheep red blood cells (sRBCs) **(B)**, zymosan **(C)**, or RFP-expressing *Leishmania major* **(D)** for 4hr and imaged by live cell confocal microscopy. Scale bars = 5 μm. **(E-G)** RawKb.APOL7C::mCherry cells were transfected with the PtdIns(3)P biosensor p40PX::GFP and challenged with OVA-beads for 4hr befor confocal imaging. Scale bar = 5 μm. Plotted are the total number of p40PX::GFP^+^ phagosomes **(F)** or the number of phagosomes that are positive for both APOL7C::mCherry and p40PX::GFP **(G)**. Each dot represents a field-of-view containing 5-20 cells (*n* = 3 independent experiments). Significance was determined using a student’s t test. **(H-J)** RawKb.APOL7C::mCherry cells were challenged with OVA-beads for 4hr before staining for LAMP1^+^ phagosomes **(F)** or the number of phagosomes that are positive for both APOL7C::mCherry and LAMP1 **(G)**. Each dot represents a field-of-view containing 5-20 cells (*n* = 3 independent experiments). Significance was determined using a student’s t test. n.s. = no significance; \**P* ≤ 0.05; \*\**P* ≤ 0.01; \*\*\*\**P* ≤ 0.0001.

**Supplementary figure 4.**
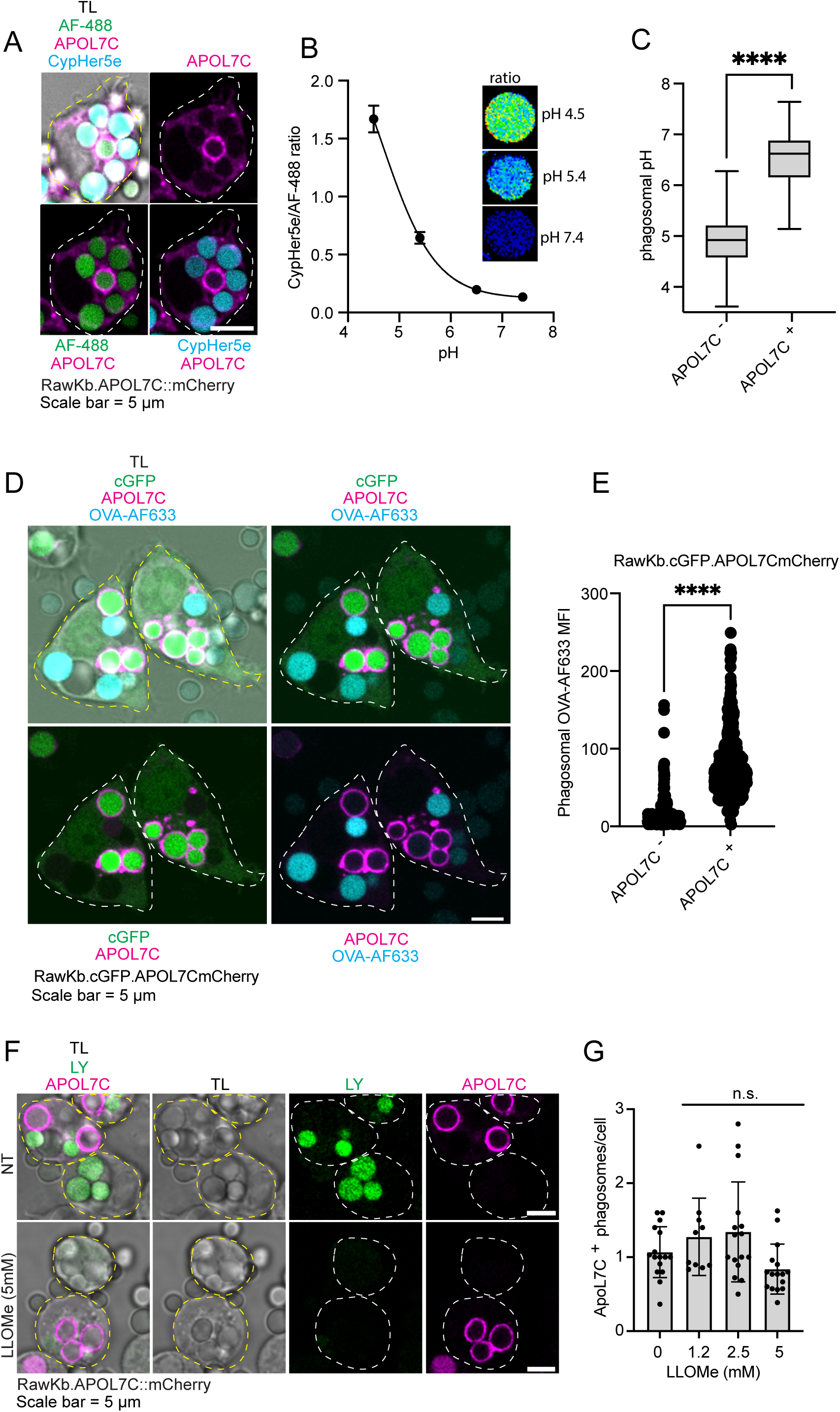
APOL7C is recruited to phagosomes independently of damage and APOL7C^+^ phagosomes are permeable to H^+^, GFP and fluorescent ovalbumin. Related to Figure 3. **(A)** RawKb.APOL7C::mCherry cells were challenged with OVA-beads labelled with Alexa Fluor 488 and the pH-sensitive fluorophore CypHer5e. They were imaged by live cell confocal microscopy 4hr after adding the beads. Scale bar = 5 μm. **(B)** A pH calibration was performed by bathing the OVA-beads from **(A)** in K^+^-rich buffers at the indicated pH. Each dot represents mean ± s.e.m. of cypHer5e/AF488 ratio of 20 single beads (*n* = 20). **(C)** Plotted is the pH of individual APOL7C^+^ (*n* = 14) and APOL7C^-^ (*n* = 109) phagosomes. Significance was determined using a student’s t test. **(D-E)** RawKb.cGFP.APOL7C::mCherry cells were incubated with beads labeled with Alexa Fluor 633 (AF633)-OVA for 4hr and then imaged by live cell confocal microscopy. Scale bar = 5 μm. Plotted is the MFI of AF633-OVA in individual APOL7C^+^ (*n* = 150) and APOL7C^-^ (*n* = 600) phagosomes, each dot represents an individual phagosome. Significance determined using a student’s *t* test. **(F-G)** RawKb.APOL7C::mCherry cells were incubated with OVA-beads and lucifer yellow (LY) for 1hr and then the extracellular LY was washed away. They were then left for an additional 3hr and subsequently treated with LLOMe at the indicated concentration. Scale bar = 5 μm. The number of APOL7C^+^ phagosomes per cell is shown, each dot represents an individual field of view containing 5-20 cells and the data is from 3 individual experiments (*n* = 3). Significance determined using a student’s *t* test. n.s. = no significance; \*\*\*\**P* ≤ 0.0001.

**Supplementary Figure 5.**
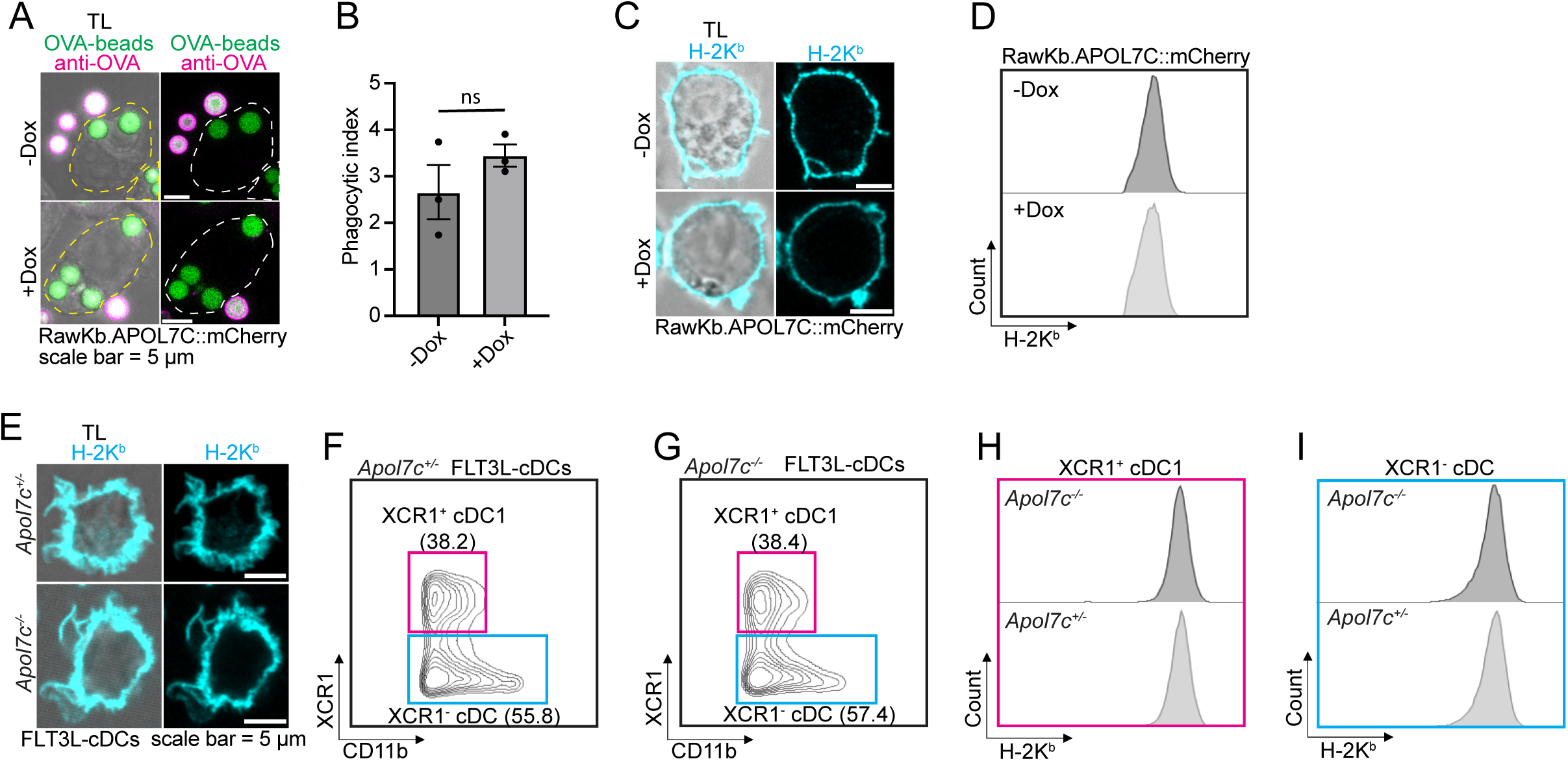
Phagocytic index for doxycycline-treated RawKb.APOL7C::mCherry cells and H-2K^b^ surface expression for RawKb.APOL7C::mCherry cells and FLT3L-cDCs. Related to Figure 4. **(A)** RawKb.APOL7C::mCherry cells were incubated with OVA-beads for 4 hours before they were fixed and stained for uninternalized beads with an anti-OVA antibody. Images were acquired by confocal microscopy. Scale bar = 5 μm. **(B)** The number of internalized OVA-beads were counted from the micrographs shown in **(A)**. Plotted as mean ± s.e.m. Each dot represents a field-of-view containing 5-20 cells (*n* = 3 independent experiments). Significance was determined using a Kolmogorov-Smirnov test. **(C)** Confocal microscopy of RawKb.APOL7C::mCherry cells stained for H-2K^b^. Scale bar = 5 μm. **(D)** Flow cytometry of RawKb.APOL7C::mCherry cells stained for H-2K^b^. Representative plot of 3 independent experiments (*n* = 3). **(E)** FLT3L-cDCs were fixed and stained for H-2K^b^. Scale bar = 5 μm. **(F-I)** Representative flow cytometry plots of FLT3L-cDCs from *Apol7c^+/-^* and *Apol7c^-/-^* mice. XCR1^+^ cDC1s were gated as CD45^+^, B220^-^, MHC-II^hi^, CD11c^hi^, CD11b^-^, XCR1^+^. XCR1^-^ cDCs were gated as CD45^+^, B220^-^, MHC-II^hi^, CD11c^hi^, XCR1^-^. H-2K^b^ histograms are derived from the gates depicted in **(F-G)**. n.s. = no significance.

**Supplementary Figure 6.**
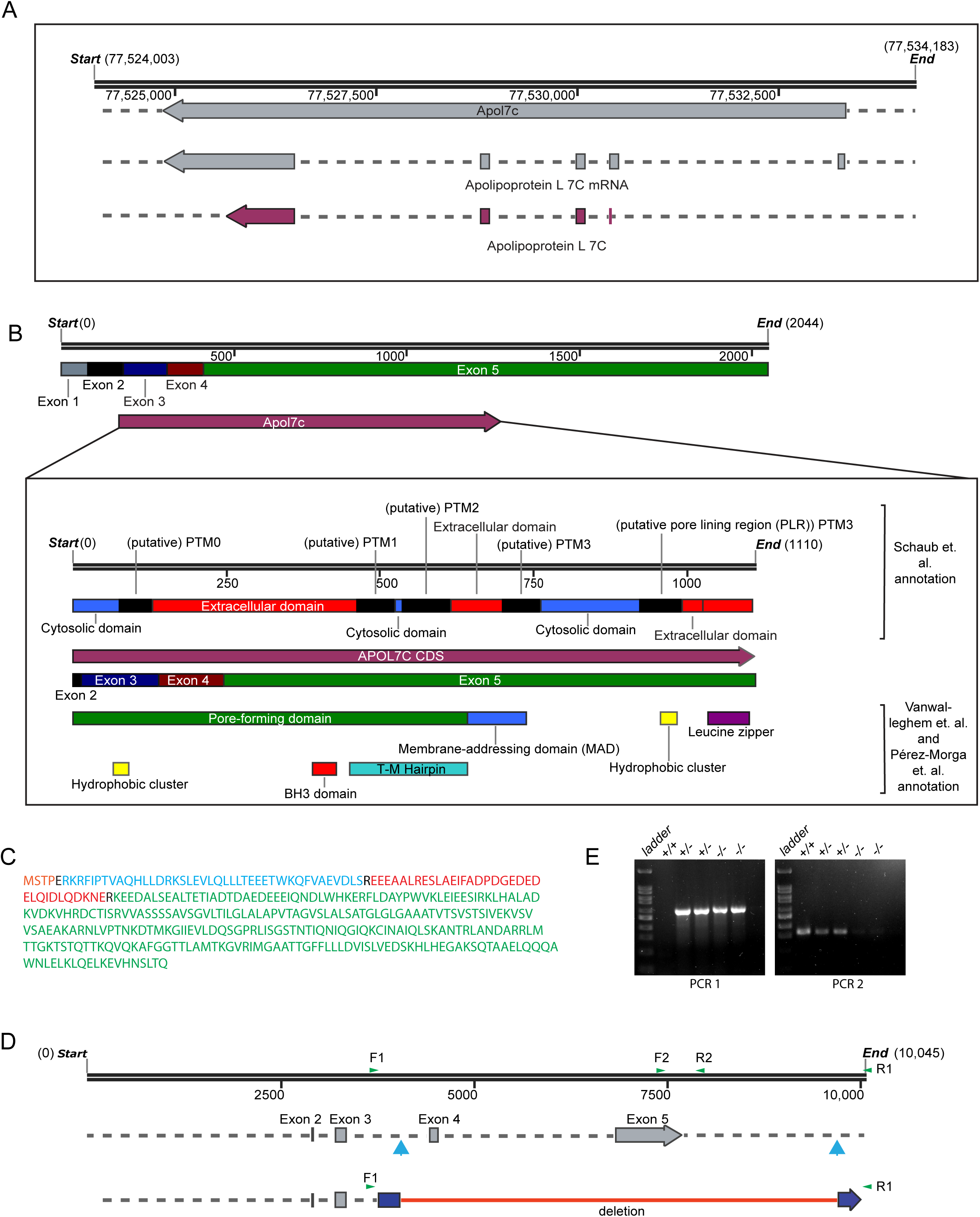
Knock-out strategy and genotyping for *Apol7c^-/-^* mouse. Related to Figure 4. **(A)** Structure of *Apol7c* gene and transcripts from NCBI. **(B)** *Apol7c* consensus coding sequence (CCDS) showing individual exons. Also shown is the domain structure of APOL7C as characterized by either Schaub *et al* or Vanwalleghem *et al* and Perez-Morga *et al*. **(C)** Protein sequence of APOL7C with alternating colors indicating contributions of individual exons (orange = exon 2, blue = exon 3, red = exon 4, green = exon 5). **(D)** Light blue arrows indicate the gRNA cut sites and green arrows indicate the primer sets used for genotyping. Primers F1 and R1 were used to amplify the deletion site and the amplicon was TOPO cloned and sequenced. The sequenced region is shown in dark blue and the deleted region is shown by the red line. **(E)** Shown is a typical genotyping result from wild-type, heterozygous and knock-out mice. PCR 1 uses primers F1 and R1 and PCR2 uses primers F2 and R2 shown in **(D)**.

**Supplementary Figure 7.**
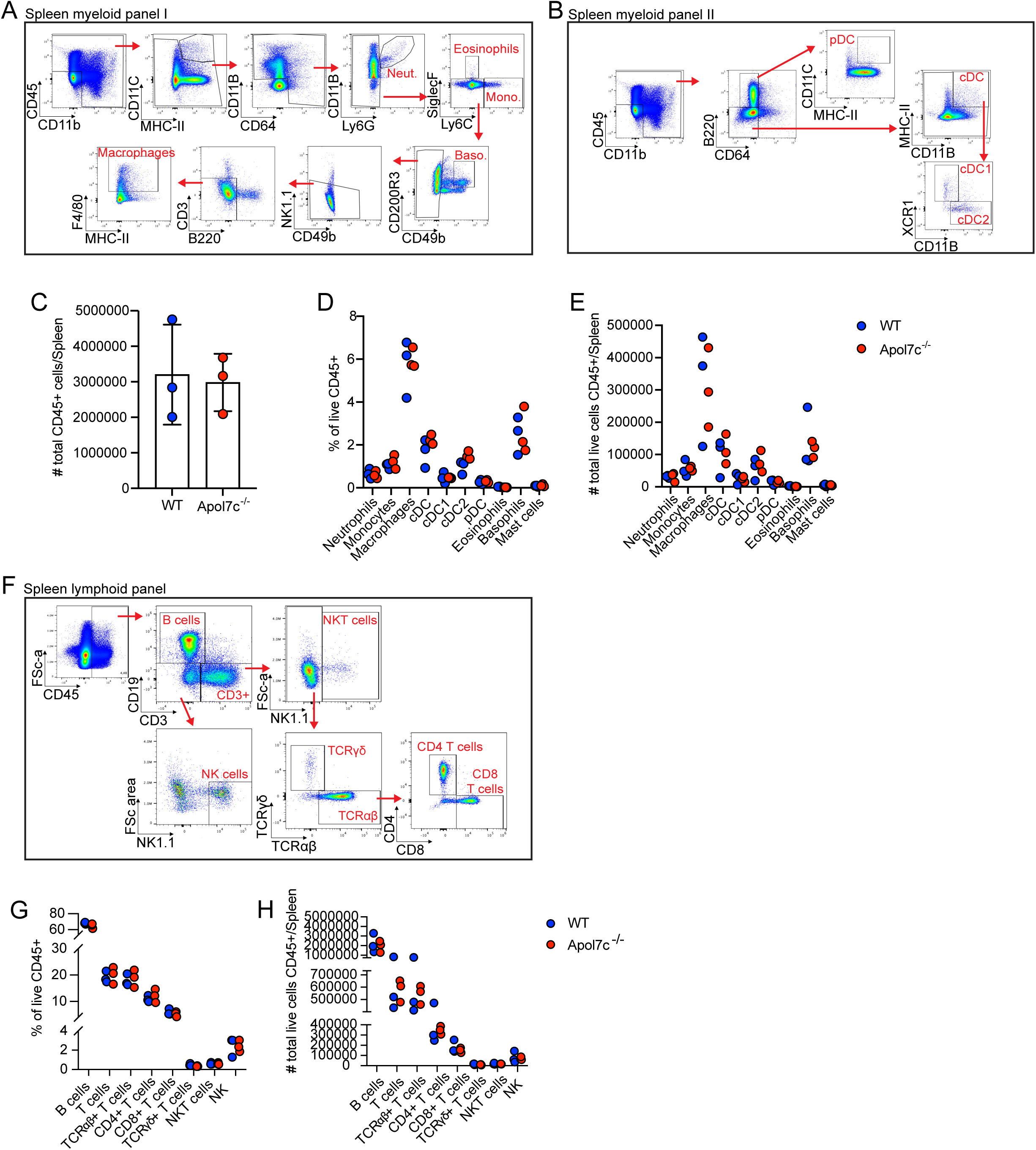
*Apol7c^-/-^* mice exhibit a normal immune profile. Related to Figure 4. **(A-B)** Flow cytometry gating strategy for myeloid cell profiling from spleen cells for both *Apol7c*^+/-^ (n = 3) and *Apol7c^-/-^* (n = 3) mice. **(C-E)** Flow cytometric analysis of the indicated immune cell populations. **(F)** Flow cytometry gating strategy for lymphoid cell profiling from spleen cells for both *Apol7c*^+/-^ (n = 3) and *Apol7c^-/-^* (n = 3) mice. **(G-H)** Flow cytometric analysis of the indicated immune cell populations.

**Supplementary Figure 8.**
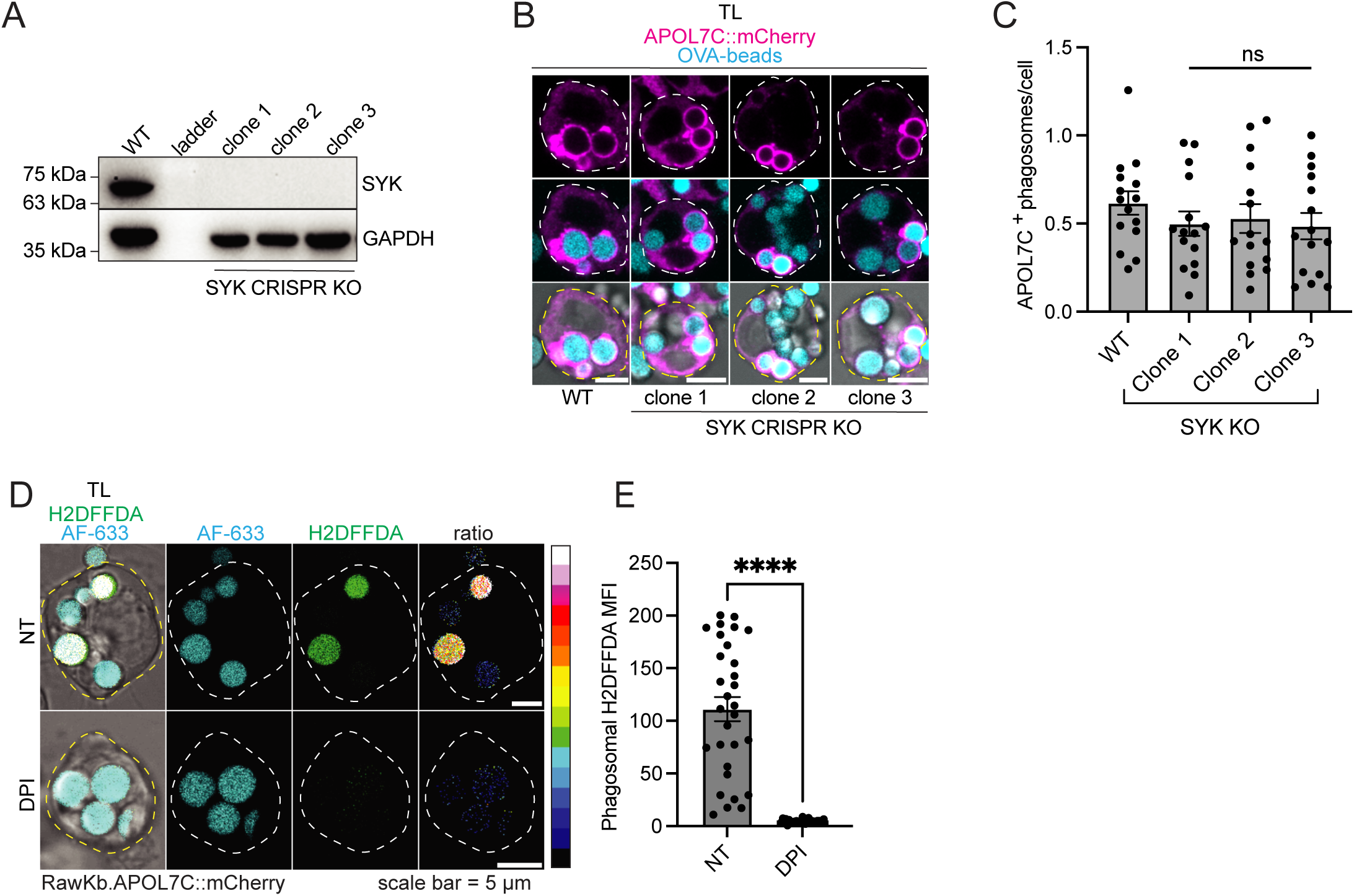
Recruitment of APOL7C::mCherry to phagosomes in SYK CRISPR knockout cells and validation of H2DFFDA as a probe for measuring NADPH oxidase activity in phagosomes. Related to Figure 5. **(A)** Western blot showing knockout of SYK in RawKb.APOL7C::mCherry cells. **(B-C)** RawKb.APOL7C::mCherry wild-type (WT) and SYK CRISPR KO cells were treated with doxycycline (DOX) overnight and then challenged with OVA-beads for 4hr. Scale bar = 5 μm. The number of APOL7C^+^ phagosomes per cell is plotted as mean ± s.e.m. Each dot represents a field-of-view containing 5-20 cells (*n* = 5 independent acquisitions). Significance determined using student’s *t* test. (**D-E**) RawKb.APOL7C::mCherry cells were incubated with Alexa Fluor 633 (AF633)- and H2DFFDA-labeled OVA-beads for 4hr in the presence of absence of DPI as indicated. Cells were then imaged by live cell confocal microscopy. Scale bar = 5 μm. Plotted is the mean ± s.e.m. of the H2DFFDA MFI in individual phagosomes (*n* = 30). Significance was determined using a Kolmogorov-Smirnov test. \*\*\*\**P* ≤ 0.0001, ns = no significance.

**Supplementary Figure 9.**
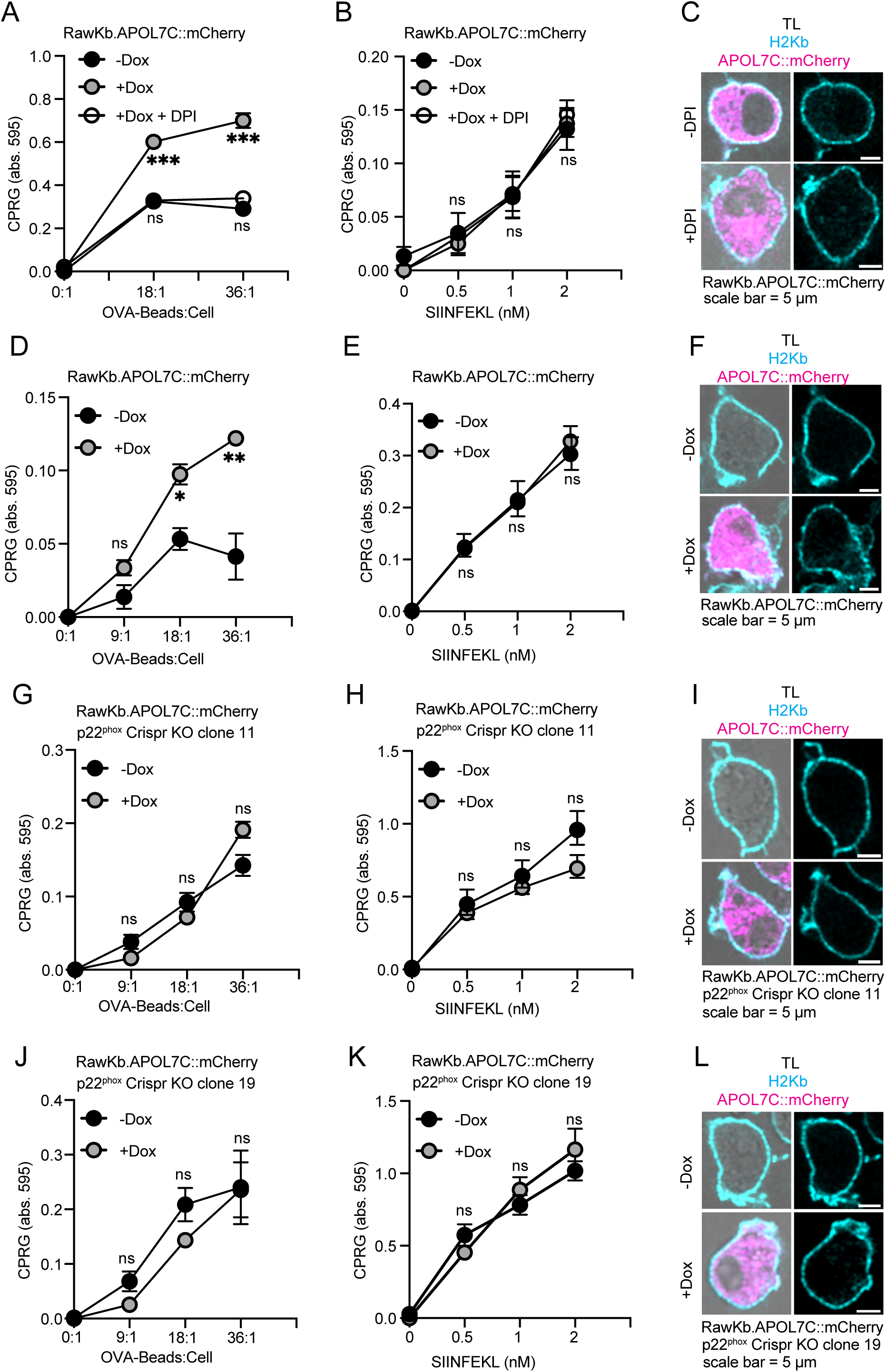
APOL7C-dependent XP requires the NADPH oxidase. Related to Figure 5. RawKb.APOL7C::mCherry **(D-F)**, RawKb.APOL7C::mCherry p22*^phox^* Crispr KO clone 11 **(G-I)**, or RawKb.APOL7C::mCherry p22*^phox^* Crispr KO clone 19 **(J-L)** were treated with or without DOX overnight and then were incubated with either OVA-beads **(A, D, G, J)** for SIINFEKL **(B, E, H, K)** in the presence or absence of DPI for 4hr. The cells were fixed and incubated overnight with B3Z cells. B3Z cells were lysed in a CPRG containing buffer. Plotted as mean ± s.d. of experimental triplicates and is representative of 3 independent experiments (*n* = 3). Significance determined using student’s *t* test for individual dilutions. The cells were also stained for H-2K^b^ and imaged by confocal microscopy **(C, F, I, L)**. Scale bar = 5 μm. n.s. = no significance; \**P* ≤ 0.05; \*\**P* ≤ 0.01; \*\*\**P* ≤ 0.001.

**Supplementary Figure 10.**
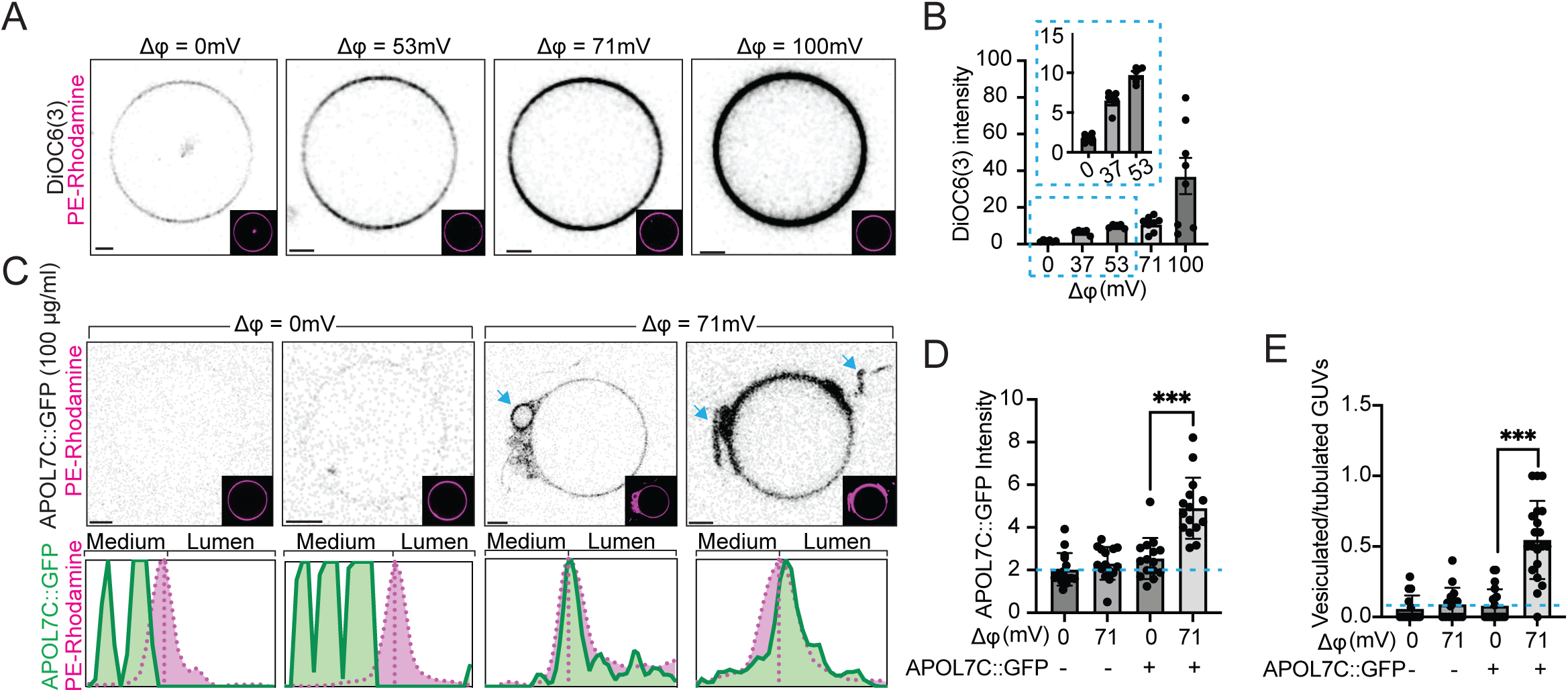
Voltage-dependent insertion of APOL7C::GFP into giant unilamellar vesicles (GUVs). Related to Figures 6 and 7. **(A-B)** GUVs containing rhodamine-labeled phosphatidylethanolamine (PE-rhodamine) and gramicidin were clamped at the indicated membrane potential by bathing in buffers of different ion composition (see methods). The membrane potential sensitive dye DiOC6(3) was added to the medium 15 mins before confocal imaging. Scale bar = 5 μm. Plotted is the mean DiOC6(3) intensity ± s.e.m. on the membrane after applying a mask from the PE-rhodamine image. Each dot represents a field of view containing 2-10 GUVs from 5-8 independent acquisitions (*n* = 5-8). Significance determined using a Kolmogorov-Smirnov test. **(C)** GUVs were prepared as in **(A)** and recombinant APOL7C::GFP (100 μg/ml) was added to the medium. Scale bar = 5 μm. **(D)** Plotted is the mean APOL7C::GFP intensity ± s.e.m. on the membrane after applying a mask from the PE-rhodamine image. Each dot represents a field of view containing 2-10 GUVs from 3 similar experiments (*n* = 3). **(E)** Plotted is the mean ± s.e.m. of vesiculated or tubulated GUVs for each of the indicated conditions. Each dot represents a field of view containing 2-10 GUVs from 3 similar experiments (*n* = 3). Significance determined by Kolmogorov-Smirnov test. \*\*\**P* ≤ 0.001.

**Supplementary video 1**. **Segmentation of APOL7C^+^ phagosome from correlative FIB-SEM. Related to** Figure 5. RawKb.APOL7C::mCherry cells were incubated with zymosan for 4hr. The cells were then imaged live by confocal microscopy to identify APOL7C^+^ phagosomes. Samples were then processed and imaged by FIB-SEM. Segmentation was performed as in methods and movie was prepared with Imaris.

**Supplementary table 1. Top 20 differentially expressed genes (DEGs) across all clusters. Related to** Figure 2. Table shows the top 20 DEGs across the clusters from scRNAseq data.

## Methods and Materials

### General materials used in this study

| Product | Catalogue number | Company |
| --- | --- | --- |
| RPMI1640 | SH30027.01 | Cytiva |
| IMDM | SH30259.02 | Cytiva |
| DMEM | SH30022.01 | Cytiva |
| DPBS | SH30028.03 | Cytiva |
| Fetal bovine serum (FBS) | 2027-06 | Corning |
| Bovine growth serum (BGS) | SH30541.03 | Cytiva |
| β-mercaptoethanol | 31350-010 | Gibco |
| Pen Strep solution | SH30010 | Cytiva |
| Puromycin | A11138-03 | Gibco |
| Hygromycin | Ant-hg-1 | Invivogen |
| Dimethyl sulfoxide (DMSO) | 0231-500ML | VWR |
| Paraformaldehyde (16 %) | 15710 | Electron Microscopy Sciences (EMS) |
| Glutaraldehyde (8 %) | 16019 | Electron Microscopy Sciences (EMS) |
| Sulforhodamine B | S1402 | Sigma Aldrich |
| In-Fusion HD Cloning Kit | 638948 | Takara |
| Prolong diamond anti-fade mountant | P36961 | Invitrogen |
| OxySens H2DFFDA, SE | 764S | Serateh Biotech |
| Alexa Fluor 488, SE | A20000 | Invitrogen |
| Alexa Fluor 555, SE | A20109 | Invitrogen |
| Alexa Fluor 633, SE | A20005 | Invitrogen |
| CypHer5e, SE | PA15401 | Cytiva |
| Zymosan, depleted | J62876 | Alfa Aesar |
| Sheep red blood cells, 10 % washed pool | 0855876 | MP Biomedicals |
| Silica beads, COOH modified | PSI-3.0COOH | Kisker Biotech |
| Polystyrene beads, COOH modified | 09850 | Polysciences Inc. |
| Fugene HD transfection reagent | 104162 | Active Motif |
| Zombie UV Live/Dead stain | 423107 | Biolegend |
| Zombie Red Live/Dead stain | 423109 | Biolegend |
| Coverslips (12 mm) | 48382 041 | VWR |
| Glass slides | 16004-388 | VWR |
| Ovalbumin | A5503-1G | Sigma Aldrich |
| 8 well live cell imaging dish | 80826 | Ibidi |
| μ-dish gridded (35 mm) | 81168 | Ibidi |
| GlutaMAX supplement | 35050-061 | Gibco |
| Glycine | BP381-5 | Fisher |
| Cyanimide | 187364-25G | Sigma Aldrich |
| Ammonium chloride | A0171-500G | Sigma Aldrich |
| R406 | S2194 | Selleckchem |
| Diphenylene iodonium (DPI) | D2926 | Sigma Aldrich |
| Liproxstatin-1 | 17730 | Cayman Chemical Company |
| ouabain | 4995-1GM | Sigma Aldrich |
| valinomycin | 10009152 | Cayman Chemical Company |
| Propidium iodide |  |  |
| 16:0-18:1 PtdCho (POPC) | 850457C | Avanti Polar Lipids Inc. |
| 16:0-18:1 PtdSer (POPS) | 840034 | Avanti Polar Lipids Inc. |
| 18:1 Liss Rhodamine PtdEth | 810150 | Avanti Polar Lipids Inc. |
| DSPtdEth-PEG(2000) Biotin | 880129C | Avanti Polar Lipids Inc. |
| Ultra-low gelling temperature agarose (Type IX-A) | Sigma Aldrich | A2576 |
| DiOC<sub>6</sub>(3) | 70009 | Biotium |
| DiBAC<sub>4</sub>(5) | 21410 | AAT Bioquest |
| Lucifer yellow | L453 | Invitrogen |
| Gramicidin A | 50845-5MG | Sigma Aldrich |
| Poly(I:C) vaccine grade | Tlrl-pic | Invivogen |
| LLOMe | L7393-500MG | Sigma Aldrich |
| CPRG | C500 | Gold Biotechnology Inc. |
| Doxycycline | J60579-14 | Alfa Aesar |
| Alt-R S.p. Cas9 Nuclease V3 | N/A | Integrated DNA Technologies (IDT) |
| Alt-R electroporation enhancer | N/A | Integrated DNA Technologies (IDT) |
| Sodium Bicarbonate solution | 17-613E | Lonza |
| DACCA SE | 506 | AAT Bioquest |
| GeneJet RNA purification kit | K0731 | ThermoFisher |
| EasySep Pan-DC enrichment kit II | 19863 | StemCell |
| qScript cDNA synthesis kit | 95047 | QuantaBio |
| iQ SYBR Green Supermix | 1708880 | BioRad |
| Osmium tetroxide, 4% solution | 19170 | Electron Microscopy Sciences |
| Potassium ferrocyanide | 470302-126 | Ward's Science |
| Durcupan ACM, Comp. A | 44611 | Sigma Aldrich |
| Durcupan, Comp. B | 44612 | Sigma Aldrich |
| Durcupan, Comp. C | 44613 | Sigma Aldrich |
| Durcupan, Comp. D | 44614 | Sigma Aldrich |
| Sodium Cacodylate Buffer, 0.2M | 11652 | Electron Microscopy Sciences |
| PELCO conductive silver paint | 16062 | Electron Microscopy Sciences |
| 5-(and-6)-Carboxy SNARF-1, acetoxymethyl ester, acetate | C1272 | ThermoFisher Scientific |
| Carbenicillin | C1389 | Sigma Aldrich |
| Overnight express instant TB medium | 71491 | Sigma Aldrich |
| Protease inhibitor cocktail powder | P2714 | Sigma Aldrich |
| DNase I | 11284932001 | Sigma Aldrich |
| Lysoszyme | L6876 | Sigma Aldrich |
| Zwittergent 3-14 | T2653 | TCI America |
| His60 Ni superflow resin | 635659 | Takara |

### Plasmids used in this study

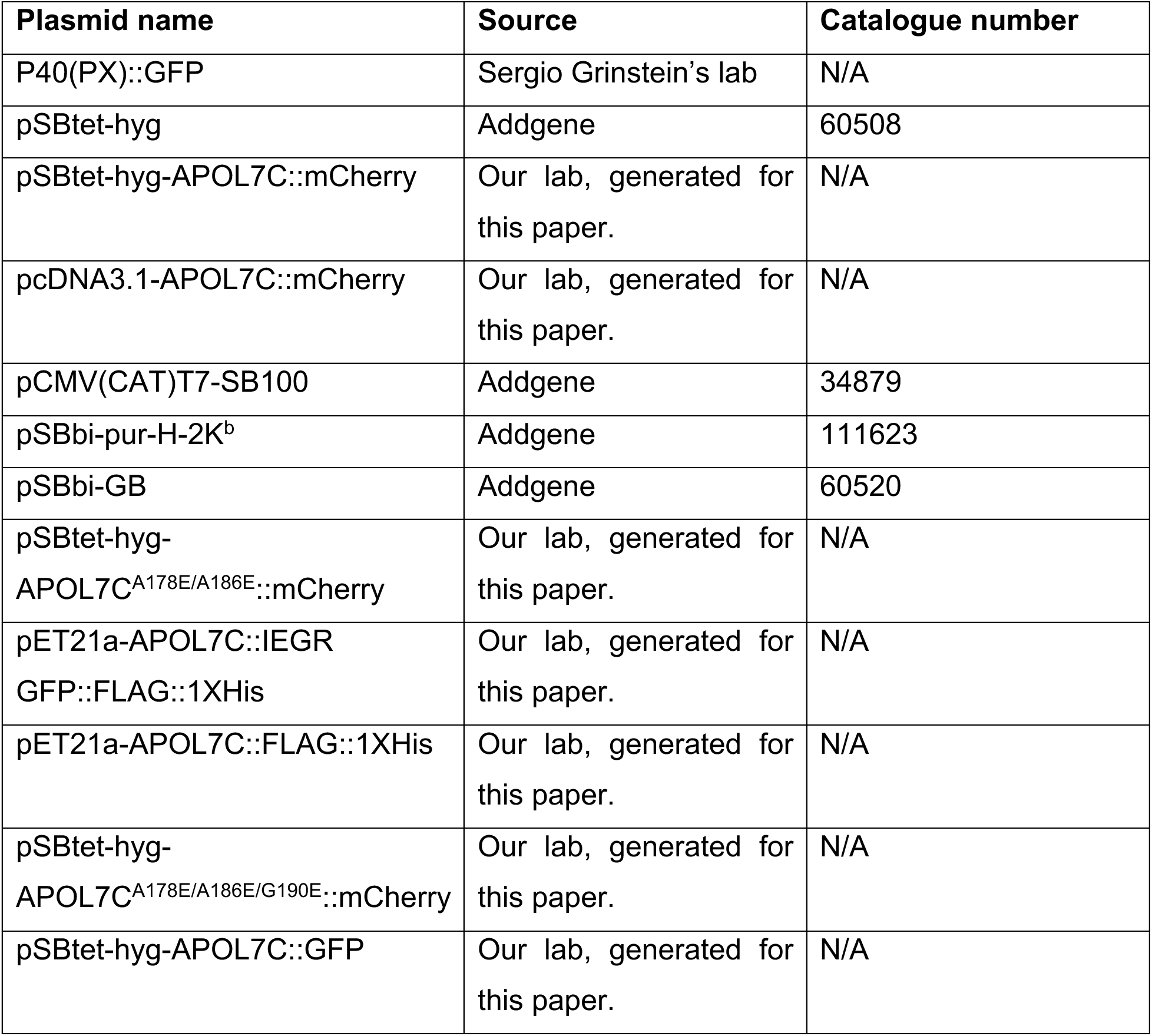

*Plasmids used in this study.*
| Plasmid name | Source | Catalogue number |
| --- | --- | --- |
| P40(PX)::GFP | Sergio Grinstein's lab | N/A |
| pSBtet-hyg | Addgene | 60508 |
| pSBtet-hyg-APOL7C::mCherry | Our lab, generated for this paper. | N/A |
| pcDNA3.1-APOL7C::mCherry | Our lab, generated for this paper. | N/A |
| pCMV(CAT)T7-SB100 | Addgene | 34879 |
| pSBbi-pur-H-2K<sup>b</sup> | Addgene | 111623 |
| pSBbi-GB | Addgene | 60520 |
| pSBtet-hyg-APOL7C<sup>A178E/A186E</sup>::mCherry | Our lab, generated for this paper. | N/A |
| pET21a-APOL7C::IEGR GFP::FLAG::1XHis | Our lab, generated for this paper. | N/A |
| pET21a-APOL7C::FLAG::1XHis | Our lab, generated for this paper. | N/A |
| pSBtet-hyg-APOL7C<sup>A178E/A186E/G190E</sup>::mCherry | Our lab, generated for this paper. | N/A |
| pSBtet-hyg-APOL7C::GFP | Our lab, generated for this paper. | N/A |

### Antibodies used in this study

#### Unconjugated primary antibodies

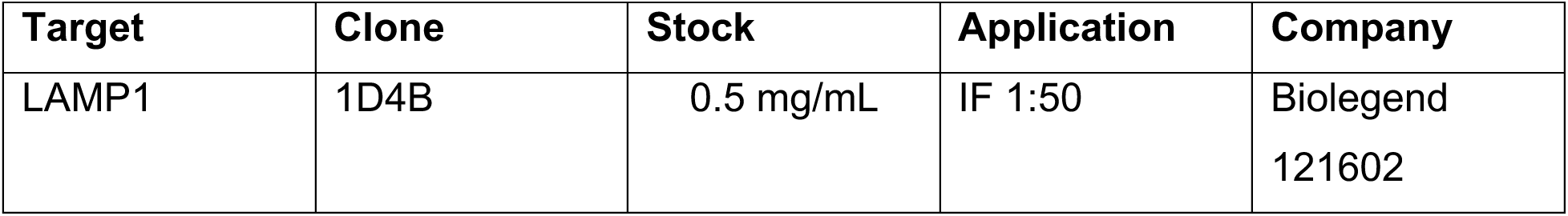
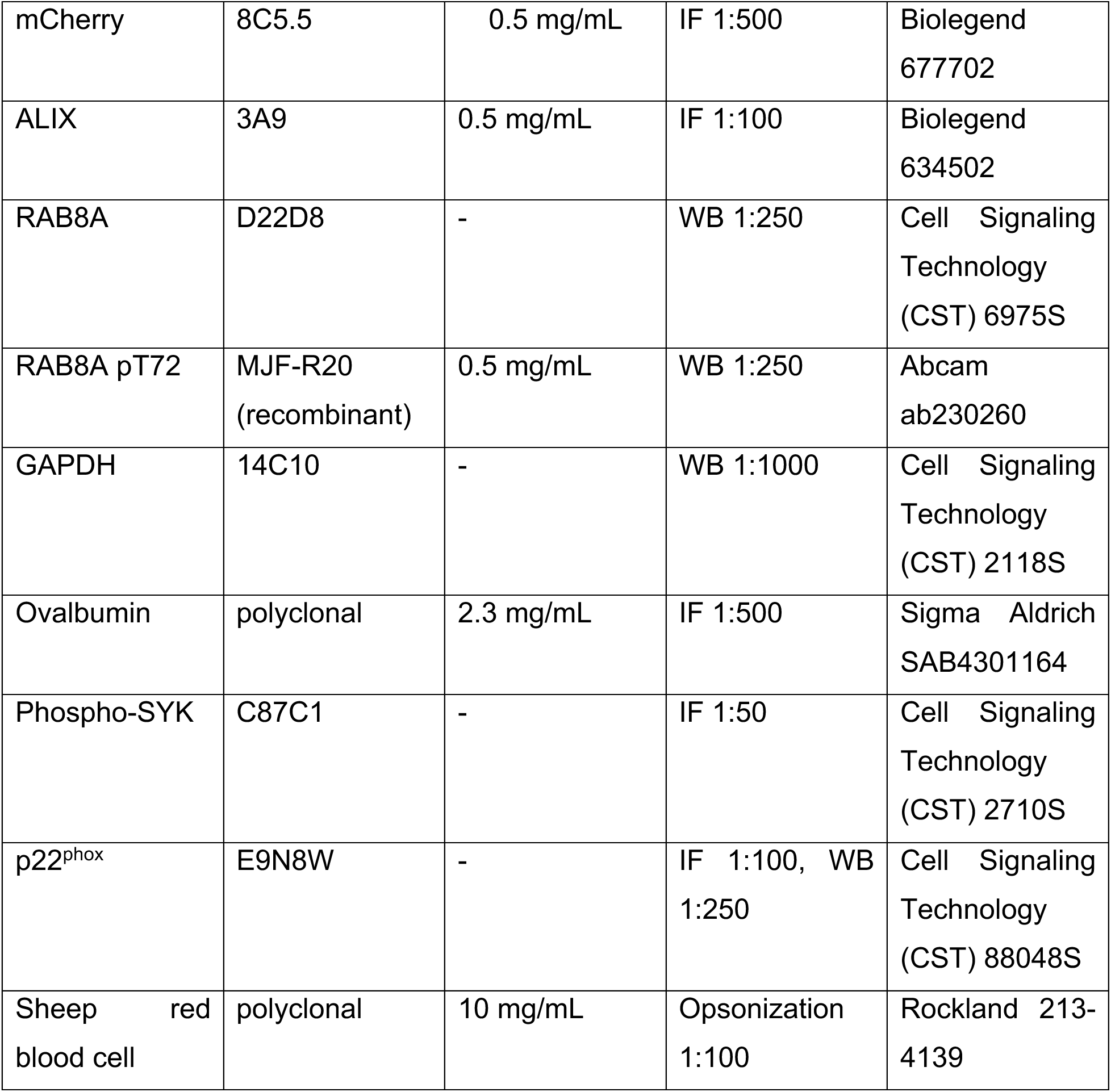

#### Conjugated primary antibodies

*Conjugated primary antibodies.*
| Target | Conjugate | Clone | Stock | Application | Company |
| --- | --- | --- | --- | --- | --- |
| Galectin 3 | AF 647 | M3-38 | 0.5 mg/mL | IF 1:500 | Biolegend<br>125408 |
| B220 | Various | RA3-6B2 | 0.2 mg/mL | FC 1:50 | Biolegend<br>103245 |
| MHC II | Various | M5/114.15.2 | 0.2 mg/mL | FC 1:300 | Biolegend<br>107623 |
| CD11c | Various | N418 | 0.5 mg/mL | FC 1:200 | Biolegend<br>117305 |
| XCR1 | Various | ZET | 0.5 mg/mL | FC 1:50 | Biolegend<br>148213 |
| CD11b | Various | M1/70 | 0.2 mg/mL | FC 1:80 | Biolegend<br>101211 |
| CD8a | Various | 53-6.7 | 0.2 mg/mL | FC 1:80 | Biolegend<br>100707 |
| CD19 | Various | 6D5 | 0.03 mg/mL | FC 1:20 | Biolegend<br>115543 |
| CD3 | Various | 17A2 | 0.5 mg/mL | FC 1:200 | Biolegend<br>100215 |
| H-2K<sup>b</sup> | AF 647 | AF6-88.5 | 0.5 mg/mL | IF 1:100, FC<br>1:50 | Biolegend<br>116511 |

#### Conjugated secondary antibodies

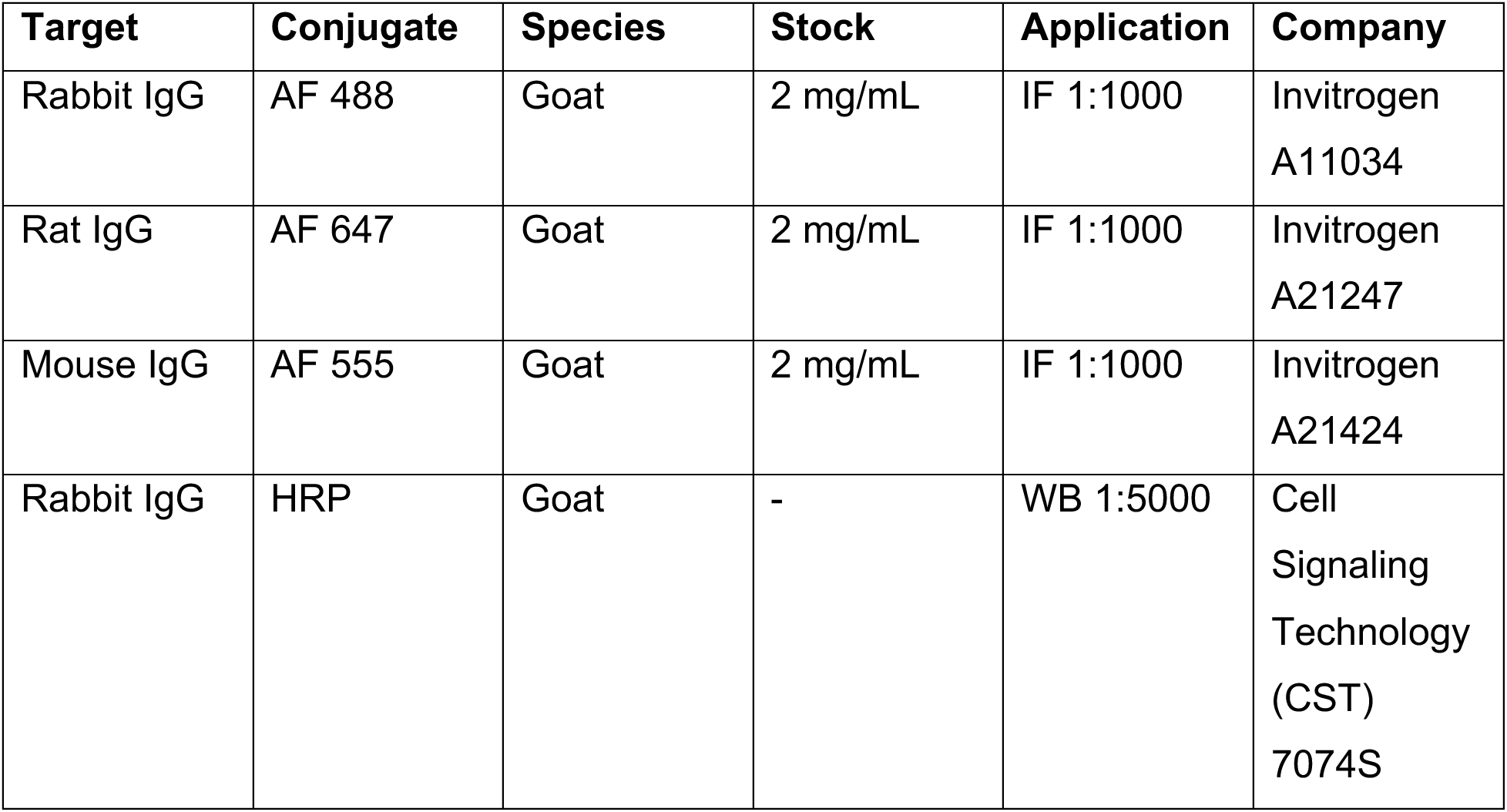
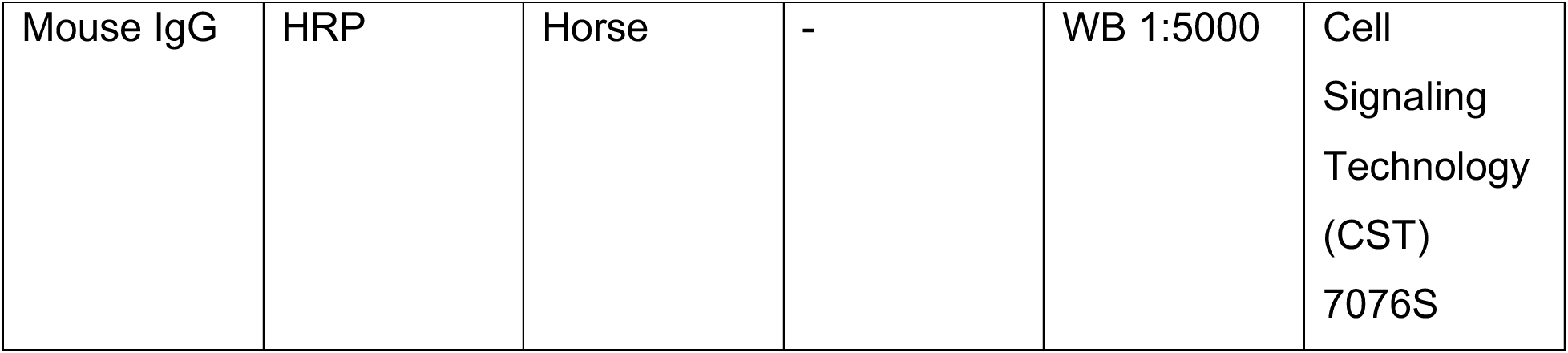

### Cells

RPMI 1640 supplemented with 10% heat-inactivated bovine growth serum (HI-BGS), penicillin, and streptomycin was used to grow Raw264.7 and all derivatives of Raw264.7 cells. Raw264.7 cells were provided by Dr. Robin Yates. RPMI 1640 supplemented with 10% HI-BGS, penicillin, streptomycin and β-mercaptoethanol was used to grow B3Z cells. B3Z cells were obtained from Dr. Albert Descoteaux. IMDM supplemented with 10% heat-inactivated fetal bovine serum (FBS), penicillin, streptomycin, glutaMAX, sodium bicarbonate and HEPES was used to grow MuTuDC1940 cells. MuTuDC1940 cells were purchased from Applied Biological Materials Inc. (ABM).

FLT3L-cDCs were generated as described by Mayer *et al*^86^. Specifically, C57BL/6 bone marrow derived cells were cultured in IMDM supplemented with 10% heat-inactivated fetal bovine serum (FBS), penicillin, streptomycin, glutaMAX, sodium bicarbonate, HEPES, 1.5 ng/mL GM-CSF and 60 ng/mL FLT3-L for 12-14 days. The culture medium was topped up at day 5 and cells were re-plated with fresh cytokines on day 9 and used between days 12-14.

### Mice

C57BL/6J, and *Batf3^-/-^* mice were obtained from Jackson. *Apol7c^-/-^* mice were generated by GemPharmatech llc by microinjecting precomplexed recombinant Cas9 and sgRNAs that targeted sequences flanking exons 4 and 5 of *Apol7c* into fertilized eggs of C57Bl/6J mice. Fertilized eggs were transplanted to obtain positive F0 mice which were confirmed by PCR and sequencing that checked for deletion of exons 4 and 5. Stable F1 generation mice were obtained by breeding the F0 mice with C57Bl/6J mice. F1 heterozygotes were shipped to the University of Calgary. Primers used for genotyping were:

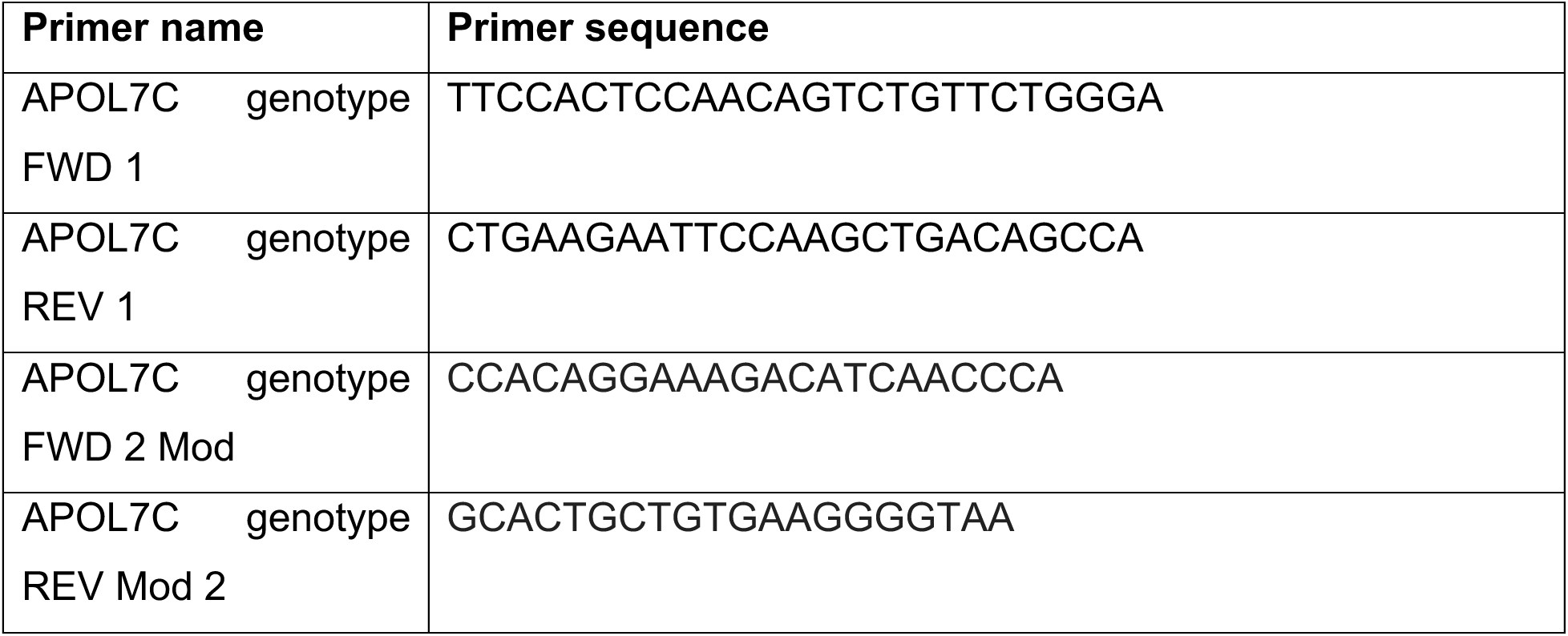

All mice were bred at the University of Calgary Foothills Campus mouse facility under specific pathogen-free conditions. Mice were used at 6-8 weeks of age and littermates were randomly assigned to experimental groups. All mouse experiments were performed in accordance with national and institutional guidelines for animal care under specific protocols that underwent institutional review.

### Isolation of splenic dendritic cells

The EasySep Mouse Pan-DC Enrichment kit was used to isolate splenic dendritic cells in accordance with the manufacturer’s instructions. The mouse was euthanized with CO_2_, followed by cervical dislocation. The spleen was harvested and then crushed in a 70 μm strainer. The splenic cells were treated with an NH_4_Cl red blood lysis buffer for 10 minutes and then spun down at 300XG for 5 minutes before being resuspended in 1ml of EasySep buffer. Next, 50 μl of the enrichment cocktail was added to the sample and incubated for 5 minutes, followed by the addition of 40 μl of RapidSphere beads for another 5 minutes. The cell suspension was then brought up to a volume of 2.5ml with EasySep buffer and transferred to a magnet for 3 minutes, resulting in collection of the pan-DC enriched supernatant.

### Generation of stable Raw264.7 and MuTuDC1940 cell lines

Stable Raw264.7 cells stably expressing H-2K^b^, APOL7C::mCherry, APOL7C^A178E/A186E^::mCherry or APOL7C^A178E/A186E/G190E^::mCherry and MuTuDC1940 cells stably expressing APOL7C::mCherry were generated using the Sleeping Beauty transposon system. H-2K^b^ in the pSBbi-puromycin vector was obtained from Addgene (deposited by Jonathan Yewdell). APOL7C::mCherry was synthesized by BioBasic in a pcDNA3.1(+) mammalian expression vector. It was subsequently cloned into the pSBtet-hygromycin vector (obtained from Addgene) using the Takara InFusion cloning system using the following primers:

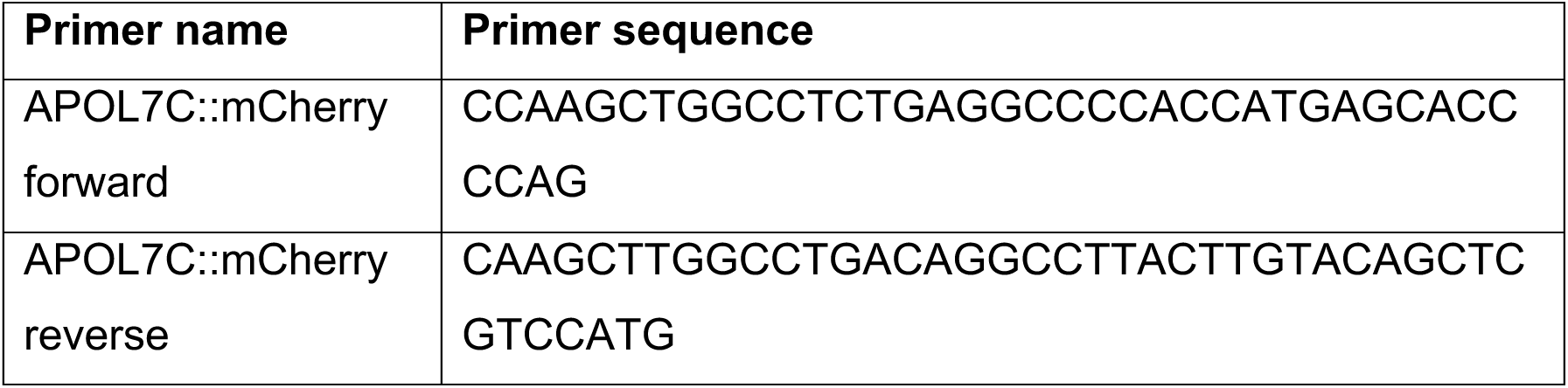

APOL7C^A178E/A186E^::mCherry or APOL7C^A178E/A186E/G190E^::mCherry in the pSBtet-hygromycin vector were generated by site-directed mutagenesis using the following primers:

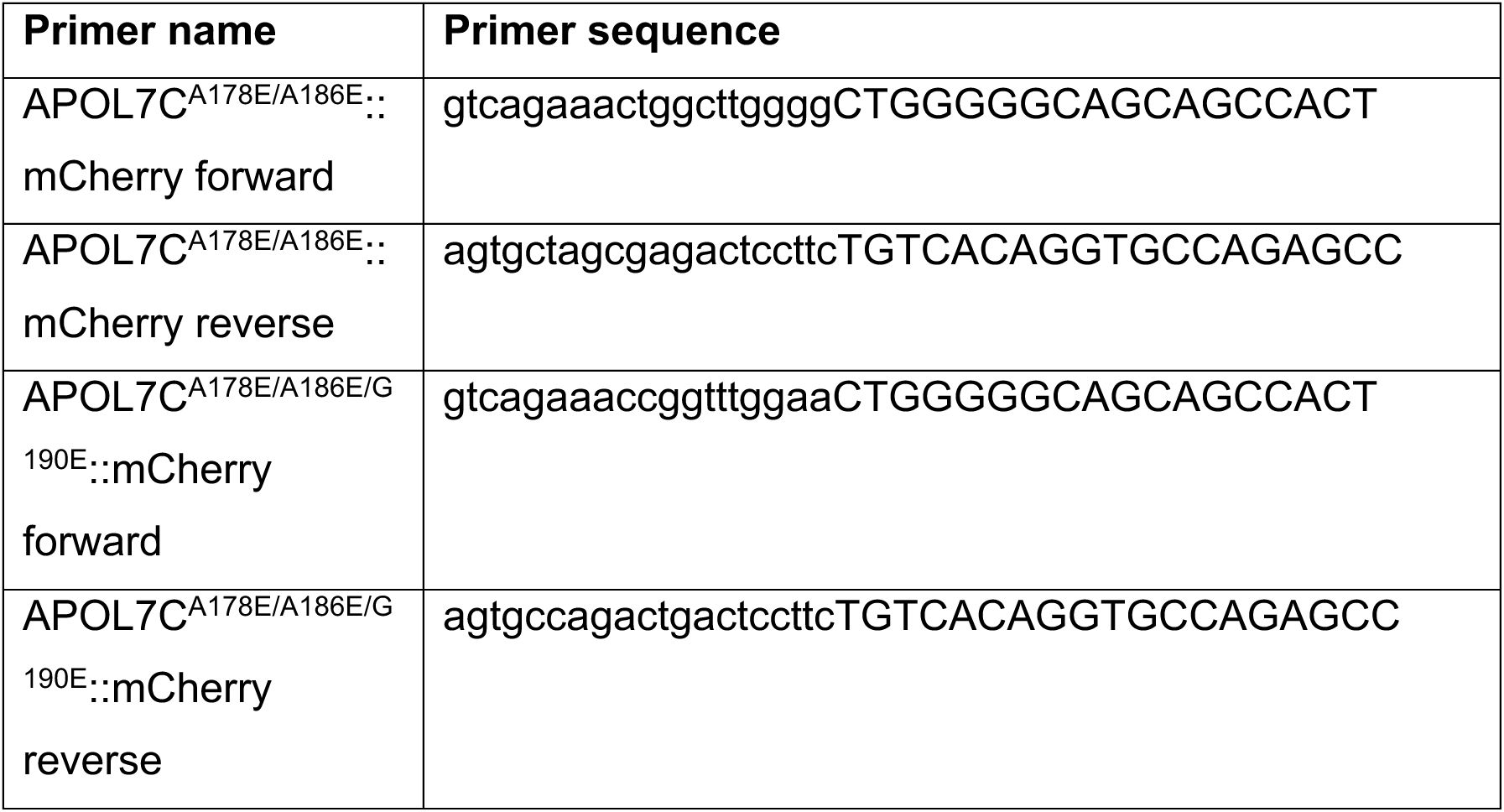

APOL7C::GFP was synthesized by BioBasic and cloned into the pET21a vector. APOL7C::GFP was then subcloned into pSBtet-hyg using the Takara InFusion system and the following primers:

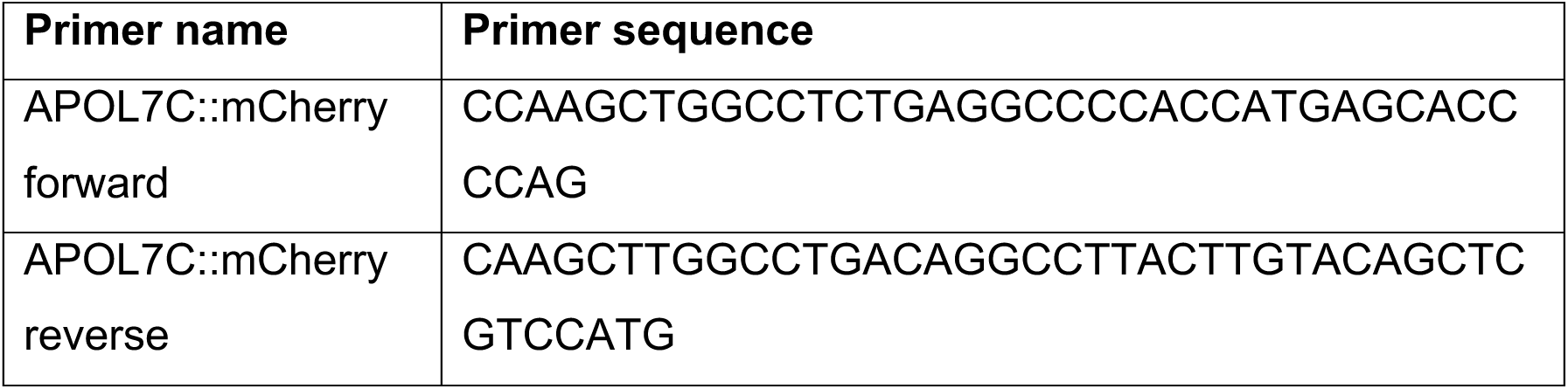

For the generation of stable cell lines, cells were transfected using the Neon Transfection system with either pSBbi-GB, pSBbi-puromycin-H-2K^b^, pSBtet-hygromycin-APOL7C::mCherry, pSBtet-hygromycin-APOL7C::GFP, pSBtet-hygromycin-APOL7C^A178E/A186E^::mCherry or pSBtet-hygromycin-APOL7C^A178E/A186E/G190E^::mCherry along with the SB100X transposase in the pCMV(CAT)T7 vector (Addgene). Transfected cells were plated into 6-well plates containing antibiotic free DMEM medium. Twenty-four hours post-transfection, the selection antibiotic was added to the medium at the concentration below:

| Selection antibiotic | Concentration |
| --- | --- |
| Puromycin | 1 µg/mL |
| Hygromycin | 300 µg/mL |

Cells were grown for 5-7 days in the selection antibiotic and then sorted for equal expression of surface H-2K^b^.

### Generation of p22^phox^ knockout RawKb.APOL7C::mCherry cell lines

RawKb.APOL7C::mCherry cells were transfected with two synthetic guide RNAs (sgRNAs) targeting p22*^phox^* precomplexed with recombinant Alt-R S.p. Cas9 nuclease V3 (IDT) using the Neon transfection system.

| sgRNA | Sequence |
| --- | --- |
| Murine CYBA sgRNA 1 | mG*mG*mC*rCrArUrGrUrGrGrGrCrCrArArCrGrArArCrGr<br>UrUrUrUrArGrArGrCrUrArGrArArArUrArGrCrArArGrUrUrA<br>rArArArUrArArGrGrCrUrArGrUrCrCrGrUrUrArUrCrArArCr<br>UrUrGrArArArArGrUrGrGrCrArCrCrGrArGrUrCrGrGrUr<br>GrCmU*mU*mU*rU |
| Murine CYBA sgRNA 2 | mU*mG*mG*rCrCrArArGrCrArGrArCrGrGrUrCrCrCrCrGr<br>UrUrUrUrArGrArGrCrUrArGrArArArUrArGrCrArArGrUrUrA<br>rArArArUrArArGrGrCrUrArGrUrCrCrGrUrUrArUrCrArArCr<br>UrUrGrArArArArGrUrGrGrCrArCrCrGrArGrUrCrGrGrUr<br>GrCmU*mU*mU*rU |

Twenty-four hours after transfection the cells single cell cloned and several weeks later the colonies were screened by western blot for p22*^phox^* expression.

### RT-qPCR for APOL7C expression

The mRNA was isolated using the Genejet RNA purification kit in accordance with the manufacturer’s instructions. To briefly summarize, cells were lysed in lysis buffer for 1 minute, followed by supplementation with 100% ethanol and vortexing for 10 seconds. The mixture was then transferred into a purification column at room temperature and spun down at 12,000 rpm for 1 minute at 4°C, and the supernatant was discarded. The column was washed with wash buffer A and B before being transferred into a new collection tube. Finally, 100 μl of nuclease-free water was added to the purification column, and it was spun down at 12,000 rpm for 1 minute to collect the flow-through. The cDNA was prepared using the QuantaBio qScript cDNA synthesis kit according to manufacturer’s instructions. To summarize, 0.1 μg of mRNA was combined with 1 μl of reverse transcriptase, 4 μl of reaction mix, and nuclease-free water was added to reach a final volume of 20 μl. The PCR reaction was run on a BioRad iQ5 thermocycler using iQ SYBR Green Supermix according to manufacturer’s instructions. Gene expression was calculated relative 18S rRNA. The primers used for APOL7C expression were:

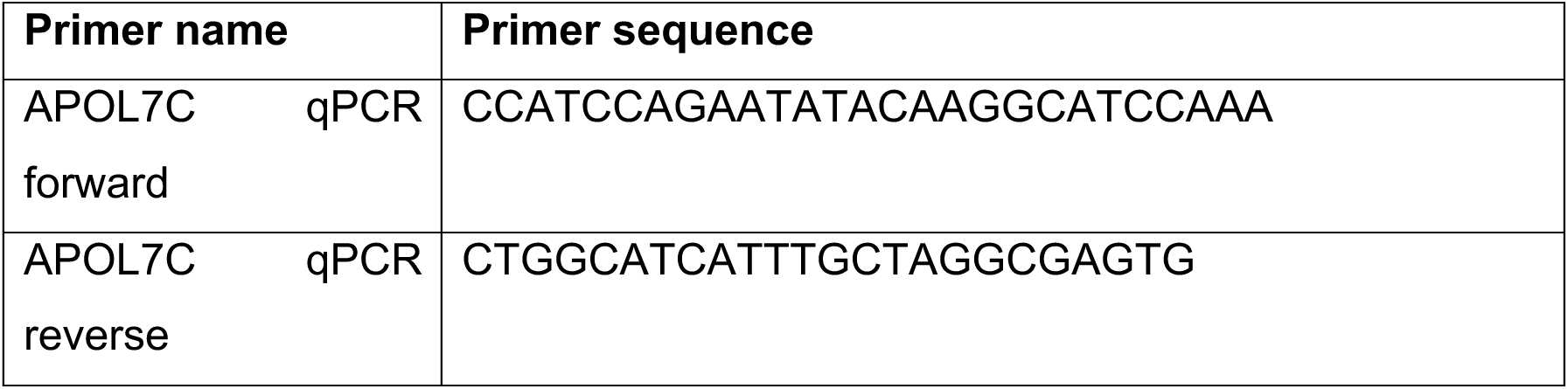

### Preparation of OVA-beads

50 mg of 3 μm carboxylate-modified silica beads were washed 3X with 1 mL of PBS in low binding 1.5 mL tubes. Beads were then resuspended in 30 mg/mL cyanimide in PBS. Beads were rotated for 15 minutes in the cyanimide solution. The beads were then washed 3X with 1 mL of 0.1 M sodium borate (pH 8.0) buffer. Beads were resuspended in 1 mL of 0.1 M sodium borate (pH 8.0) buffer containing 3 mg/mL endotoxin-free ovalbumin. Beads were rotated for 2 hours at room temperature and then washed 3X with 1 mL of PBS to remove excess ovalbumin and resuspended in 1 mL of PBS. 5 μL of a 2 mg/mL stock of a succinimidyl ester derivative of the desired fluorophore was then added to the tube and the beads were rotated for 1 hour at room temperature. Beads were then washed 3X with 1 mL of PBS to remove excess dye and resuspended in 1 mL of PBS containing 0.2M glycine. The beads were rotated at 4°C overnight and then washed 3X with 1 mL of PBS.

### Preparation of Staphylococcus aureus and infection

*Staphylococcus aureus* strain MW2 was obtained from NARSA (Network on Antimicrobial Resistance in *Staphylococcus aureus*). This strain was transformed with pCM29 to constitutively express high levels of GFP^87^. Staphylococci were grown in brain heart infusion at 37°C while shaking in the presence of chloramphenicol (10 µg/ml) for overnight maintenance of the plasmids. For infection experiments, *S. aureus* strains were sub-cultured without antibiotics until exponential phase (OD_660nm_, 1.0) washed with saline once, resuspended in saline, and added to the cells. *S. aureus* expressing GFP was adjusted to a concentration of 4.4 × 10^8^ in serum-free RPMI. They were then added to the RawKb.APOL7C::mCherry cells at an MOI of 10 bacteria to each cell (10:1). After a 1-minute centrifugation at 300 XG, the infection was left to proceed for 4 hours at 37°C and 5% CO_2_. Cells were then fixed in 4% PFA and imaged by confocal microscopy on a Leica SP8 equipped with a HC PL APO 63×/1.40 Oil CS2.

### Preparation of Leishmania major and infection

*L. major* expressing RFP (kind gift of Dr. Nathan C. Peters) was grown to log phase and metacyclic promastigotes were isolated by centrifugation on a ficoll gradient. Parasites were incubated with RawKb.APOL7C::GFP cells at an MOI of 4 parasites to each cell (4:1). They were placed at 34°C with 5% CO_2_ for 6 hours. Cells were then fixed in 4% PFA and imaged by confocal microscopy (Leica SP8 equipped with a HC PL APO 63×/1.40 Oil CS2).

### Preparation of zymosan-AF488

Zymosan (1 mg/mL) was incubated with a succinimidyl ester derivative of Alexa Fluor 488 (AF488-SE) in 0.1 M sodium borate (pH 8.0) buffer at a ratio of 0.1 mg of AF488-SE to 1 mg of zymosan. The reaction was left to proceed on a rotator for 1 hour at room temperature. The zymosan was then washed 6X with PBS and stored in PBS at 1 mg/mL at -20°C until use.

### Preparation of IgG-opsonized sheep red blood cells (IgG-sRBCs)

100 μL of a 10% suspension of sRBCs was washed 3X with 1 mL of PBS. The sRBCs were opsonized with 3 μL of rabbit IgG fraction for 1 hour at room temperature. The opsonized sRBCs were then washed 3X with PBS. To fluorescently label the IgG-sRBCs, they were incubated with a fluorophore labeled anti-rabbit secondary antibody. The IgG-sRBCs were then washed 3X with PBS and used immediately.

### Clamping pH of cells

Cells were bathed in a K^+^-rich medium (143 mM KCl, 1 mM MgCl_2_, 1 mM CaCl_2_, 5 mM glucose, and 20 mM HEPES or acetic acid depending on the pH of the medium) containing 10 μg/ml nigericin. Images were acquired at least 30 minutes after the addition of the nigericin-containing medium.

### Phagocytosis assays

MuTuDCs, MuTuDC.APOL7C::mCherry, RawKb, or RawKb.APOL7C::mCherry cells were plated onto Ibidi 8-well μ-dishes and left to adhere overnight. In the case of primary splenic cDCs or FLT3L cultures the cells were plated onto 12 mm glass coverslips that were coated with anti-MHC II. The cells were then challenged with the phagocytic targets (OVA beads, zymosan-AF488, or IgG-sRBCs) for 3 hours (unless otherwise indicated) and immediately fixed with 4% PFA in PBS. PFA was then quenched with 50 mM NH_4_Cl in PBS for 5 minutes followed by 3 washes with PBS. Outside particles were labeled with an anti-OVA antibody or an anti-rabbit antibody for OVA beads and IgG-sRBCs, respectively. Next, the cells were permeabilized with 0.4% Triton X 100 in PBS for 10 minutes followed by 3 washes with PBS. Cells were blocked for 30 minutes with 2% low fat milk in PBS and then incubated with the primary antibodies for 1-2 hours are room temperature. Cells were then washed 3X with 2% low fat milk in PBS. Secondary antibodies were then added for 1 hour, followed by 3 washes. Cells were then left in PBS for imaging (μ-dishes) or were mounted onto glass slides using ProLong Diamond mounting medium. Images were then acquired on a Leica SP8 confocal microscope (equipped with a HC PL APO 63×/1.40 Oil CS2) and the number of particles internalized per cell (referred to as the phagocytic index) was determined.

### Sulforhodamine B (SRB) and Lucifer Yellow (LY) release assays

Cells were incubated with phagocytic targets in the presence of fluid-phase SRB and LY (both at 150 μg/mL) to allow for internalization of SRB and LY into the lumen of phagosomes. After 1-4 hours of phagocytosis, the excess SRB and LY was washed off with serum-free RPMI. Cells were then immediately imaged live at at 37°C, 5% CO_2_ on a Leica SP8 confocal microscope (equipped with a HC PL APO 63×/1.40 Oil CS2) and the resulting time-lapse images were analyzed in Fiji (ImageJ2, version 2.9.0/1.53t, build a33148d777). SRB or LY fluorescence was normalized to a reference fluorophore that was covalently attached to the OVA beads as above.

### Preparation of recombinant protein

Recombinant APOL7C protein was overexpressed and purified from inclusion bodies in bacteria following the method described for purifications of the structurally similar APOL1 protein by Verdi *et al*^88^ with some modifications as below. *E. coli* BL21(DE3)-pLysS was transformed with plasmid DNA encoding *Apol7c*, C-terminally tagged with the FLAG epitope (DYKDDDDK) followed by a 6x Histidine tag, synthesized by Biobasic (Markham, ON), cloned 5’-NdeI to 3’-NotI into pET21a(+) (Novagen, Madison, WI) (pET21a-APOL7C::FLAG::1XHis). A freshly streaked colony from Luria-Bertani (LB) plates, supplemented with 100 µg/ml carbenicillin was inoculated into 1 mL of liquid LB broth with 100 µg/ml carbenicillin, shaken at 37°C for 2 hours, and then sub-cultured to 200 mL of liquid Overnight Express Instant TB medium supplemented with 100 µg/ml carbenicillin and shaken for 16 hours at 37°C. Cells were pelleted at approximately 1000 x g for 10 minutes, and supernatants completely removed. Cell pellets were frozen at -70°C overnight. Pellets were thawed, resuspended in 18 mL lysis buffer (50 mM Tris pH 8.0, 1 mM EDTA, 5 mM DTT, 1 mM PMSF, supplemented with broad-spectrum protease inhibitor (Sigma, P2714) on ice. Resuspended pellet was then sonicated (10 second pulses with 10 second rests) for six minutes. Lysates were treated with DNase I and RNase I (both 5 µg/ml) and 0.25 mg/ml lysozyme for 30 minutes at room temperature with constant stirring, followed by addition of 3.5 mM MgCl_2_, 2 mM CaCl_2_, 0.5% (v/v) Triton X-100 and sodium deoxycholate for an additional 30 minutes at room temperature, then centrifuged at 26,000 x g for 30 minutes at 4°C. Supernatant was completely discarded, and the pellet resuspended in 10 mL lysis buffer containing 0.5% Triton X-100 and 0.1 M NaCl, stirred for 30 minutes at room temperature, then centrifuged at 17,050 x g for 30 minutes at 4°C. The pellet was again resuspended in lysis buffer containing 0.1 M NaCl and stirred for 30 minutes at room temperature. Sample was transferred to ice and the sonication repeated. The solution was centrifuged at 17,050 x g for 30 minutes. Supernatant was discarded, and the protein was solubilized by resuspension of the pellet in 7 mL of sterile water containing 1x Sigma P2714 protease inhibitor. The solution was stirred with the addition of 2 mL of 10% Zwittergent 3-14 for 2 minutes, followed by dropwise addition of 100 µL of 1 N NaOH and stirring for 30 seconds. The now clarified solution was immediately neutralized by addition of 1 mL of 10x TBS (0.5 M Tris pH 7.5, 1.5 M NaCl). Insoluble material was removed by centrifugation at 17,050 x g for 30 minutes at 4°C. Supernatant of this centrifugation was filtered through a 0.45 µm polyethersulfone (PES) membrane and then purified using His60 Nickel agarose Superflow Resin through a gravity-flow column at 4°C, according to the manufacturer’s recommendations.

### Giant unilamellar vesicle (GUV) assays

GUVs were prepared on agarose films overlayed on glass slides. Specifically, 1 % ultra-low gelling temperature agarose was spread over glass coverslips and dried in a 37°C incubator until the agarose layer was near invisible. Lipid mixes were prepared (according to the recipe below) in glass vials using chloroform as the solvent.

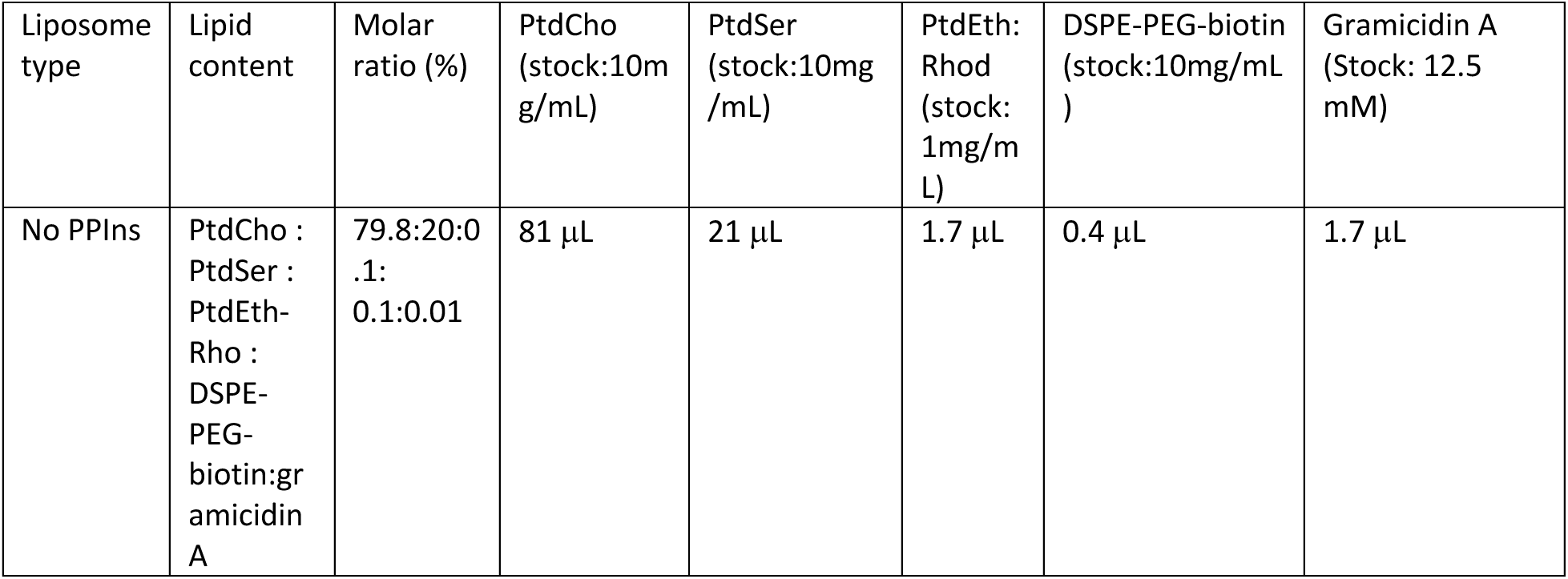

Where indicated, gramicidin A was included in the lipid suspension. Lipid suspensions were then dried under a continuous stream of argon or nitrogen gas and resuspended a small volume of chloroform (roughly 1.33 μmol of lipid in 35 μL of chloroform). The lipids were then spread over the agarose film using a glass rod taking care to make an even distribution of the lipids. The film was then dried under argon. Slides were then submerged in assay buffer (150 mM KCl, 1mM EGTA, 10 mM HEPES, pH 7.4) (hereafter K^+^-rich buffer) to allow for GUV formation. GUVs were harvested 4 hours later, stored at 4°C and were only used for 3-4 days.

### Measurement of membrane potential in GUVs

A membrane potential in the GUVs was induced as in [ref.^74,75^]. Specifically, the medium in which the GUVs were bathed was changed to a mixture of K^+^-rich buffer and a second buffer (150 mM tetraethylammonium chloride [TEAC], 1mM EGTA, 10 mM HEPES, pH 7.4) (hereafter TEAC-rich buffer). In this way the concentration of K^+^ in the bathing medium could be altered while keeping the osmolarity constant. This will impose K^+^ gradient across the GUV membrane and the resulting membrane potential can be estimated using the Nernst equation. To ensure that a membrane potential was successfully applied, the membrane potential sensitive dye 3,3’-dihexyloxacarbocyanine iodide [DiOC_6_(3)] was added at a concentration of 1 nM to the medium bathing the GUVs and imaged after 10 minutes on a confocal microsocope. Quantification of DiOC_6_(3) accumulation on GUVs was achieved through masking of the DiOC_6_(3) signal with the PtdEth:Rhod segmented signal and measuring the fluorescence intensity on the confocal images.

### Measurement of lipid peroxidation

Cells were incubated in serum-free RPMI medium containing 1 μM BODIPY 581/591 C11 for 30 minutes at 37°C. The cells were then washed 3 times with warm serum-free RPMI. OVA-beads were added for 3 hours. Uninternalized OVA beads were then washed with warm serum-free RPMI and the cells were imaged live at at 37°C, 5% CO_2_ on a Leica SP8 confocal microscope equipped with a HC PL APO 63×/1.40 Oil CS2 using a 488 nm excitation laser and emission was collected at 510 nm.

### Phagosomal membrane potential measurements

Phagosome membrane potential measurements were performed as previously done by the Grinstein lab^73^. Specifically, OVA beads with or without a DACCA label were first bathed in PBS with or without 250 nM DiBAC_4_(5). This allowed for sensitized emission FRET optimization on a Leica SP8 confocal microscope equipped with a HC PL APO 63×/1.40 Oil CS2 by acquiring images under the following configurations: donor(DACCA) exc. 405 nm and ems. 480 ± 50 nm; acceptor[DiBAC_4_(5)] exc. 552 nm and ems. 640 ± 30 nm; FRET exc. 405 nm and ems. 640 ± 30 nm. Corrected FRET (cFRET) was determined as previously described^73^. Donor and acceptor bleed-through coefficients were determined by acquiring donor alone and acceptor alone control images in all three channels described above. This correction was performed for each independent experiment. Next, cells were incubated with DACCA-labeled OVA beads in serum-free RPMI for the indicated time and then the medium was removed and replaced with fresh serum-free RPMI containing 250 nM DiBAC_4_(5) for 7 minutes. cFRET in cells was characterized by imaging as described above immediately afterwards.

### Ouabain and valinomycin treatment

Cells were incubated in serum-free RPMI containing 1 mM ouabain for 3 hours at 37°C. Next cells were washed 3 times with a K^+^- rich medium (143 mM KCl, 1 mM MgCl_2_, 1 mM CaCl_2_, 5 mM glucose, and 20 mM HEPES, pH 7.2) containing 1 mM ouabain and 1 μM valinomycin. Cells were then bathed in K^+^- rich medium containing 1 mM ouabain, 1 μM valinomycin and, where indicated, propidium iodide (PI) was included at 5 μg/mL. Imaging was then performed on live cells on a confocal microscope.

### In vitro XP assay in MuTuDCs, RawKb, and RawKb.APOL7C::mCherry

APCs were seeded at a density of 5 × 10^4^ cells per well in a 96-well U bottom plate. Where inhibitors were added, cells were pretreated with the inhibitor at the indicated concentration for 30 minutes prior to adding OVA beads. OVA beads or SIINFEKL peptide was then added at the indicated bead:cell ratio or SIINFEKL concentration for 4 hours for beads and 1 hour for SIINFEKL peptide, unless otherwise stated. Cells were then fixed with 0.008% glutaraldehyde at 4°C for 3 minutes, followed by immediate quenching with cold 0.2 M glycine in PBS for 5 minutes. Cells were then washed once with PBS and once with RPMI supplemented with 10% HI-BGS, penicillin, streptomycin and β-mercaptoethanol. Finally, 2 × 10^5^ B3Z cells in RPMI supplemented with 10% HI-BGS, penicillin, streptomycin and β-mercaptoethanol were added to each well. B3Z cells are a hybridoma line that express a T cell receptor specific for OVA(257-264) (SIINFEKL) loaded in H-2K^b^. B3Zs carry a β-galactosidase construct driven by NFAT elements from the IL-2 promoter. After 24 hours, the plates were centrifuged to pellet the B3Zs and washed 3X with PBS. B3Zs were lysed with a lysis buffer containing CPRG, the substrate for β-galactosidase, for 3-4 hours. β-galactosidase activity was measured on a plate reader (PerKin Elmer, Wallac EnVision, 2104 multilabel plate reader) capable of reading absorbance at 590 nm.

### Bulk RNAseq experiments

RNA was isolated from the MuTuDC1940 cells using the GeneJet RNA purification kit. Quality control of the preparation was performed by TapeStation RNAScreen Tape. RNASeq reads were pseudo-mapped to NCBI RefSeq transcript models using Kallisto 0.42.4^89^. Mapping counts were analyzed using Sleuth^90^, with a regression model of the binary factor treated:untreated. Differentially expressed transcripts were those that pass the aforementioned regression model with both the Wald and Likelihood Ratio Test p-values < 0.05 after Benjamini-Hochberg multiple testing correction. Transcripts were annotated using Ingenuity Pathway Analysis (Qiagen).

### Single-cell RNA-seq sample processing, library construction and sequencing

The spleen underwent the same processing steps as described for generating a single cell suspension from the spleen for pan-DC enrichment above. Next, half of the sample was enriched for pan-DCs. The remaining cells were stained with PE-anti mouse CD45 antibodies, and CD45^+^ cells were isolated using the EasySep Release PE positive selection kit, following the manufacturer’s instructions. After the isolation, an equal amount of CD45^+^ cells and pan-DCs were mixed and resuspended in 1% BSA PBS. A total of 12,000 single cells isolated from each spleen was loaded for partitioning using 10X Genomics NextGEM 3’ Gene Expression Kit (v3.1). Samples were processed according to the manufacturer’s protocol (with the PCR amplification steps run at 12X, and 14X respectively). Quality control of the final cDNA libraries was performed using TapeStation D1000 assay. Sequencing was performed using Illumina NovaSeq S2 and S1 100 cycle dual lane flow cells over multiple rounds at the UCalgary Centre for Health Genomics and Informatics (CHGI). A total of 17,027 cells were captured across each of the samples. Each sample was sequenced to approximately 15,000 reads per cell. Sequencing reads were aligned using CellRanger 3.1.0 pipeline to the standard pre-built mouse reference genome mm10 (Ensembl 93). Samples that passed alignment QC were aggregated using CellRanger aggr with between-sample normalization to ensure each sample received an equal number of mapped reads per cell in the final dataset.

### Single-cell RNA-Seq bioinformatics

Filtered feature-barcode HDF5 matrices from aggregated datasets were imported into the R package Seurat v.4.3.0 (for normalization, scaling, integration, Louvain clustering, dimensionality reduction, differential expression analysis, and visualization^91,92^. Briefly, cells with fewer than 200 UMIs, greater than 4,000 UMIs, and greater than 7% of mitochondrial reads were excluded from subsequent analysis. Cell identity was annotated based on previously published markers^59–62^.

### In vivo immunizations with OVA-coated beads and poly(I:C)

6–8 week-old mice received an intravenous injection (i.v.) of OVA-coated polyacrylamide 3.0 μm beads mixed with or without poly(I:C) (50 μg/mouse) in a total volume of 0.1 ml. 4-5 days later the mice were euthanized, spleens were removed and CD8 T cell responses were measured by staining for H-2K^b^-OVA-pentamer positive CD8 T cells.

### Coating coverslips with anti-MHC II

Glass coverslips (12 mm) were covered in a 0.1 % solution of poly-L-lysine for 30 minutes at room temperature. Coverslips were then washed 3X with PBS and incubated in 2.5 % glutaraldehyde for 15 minutes at room temperature. Coverslips were then washed 3X with PBS. A solution of 1 μg/mL of anti-MHC II in PBS was then overlayed on the coverslips for 30 minutes at room temperature. Coverslips were then washed 3 more times with PBS and then incubated overnight in 0.2 M glycine in PBS at 4°C. On the day of the experiment, coverslips were washed an additional 3 times with PBS prior to attaching cells.

### Western blots

Equal numbers of cells were plated and treated as indicated in the figure legend. Cells were then lysed directly in Laemmli buffer containing β-mercaptoethanol. Cells were then run on an SDS-PAGE, imaged and analyzed in Fiji image analysis software (ImageJ2, version 2.9.0/1.53t, build a33148d777).

### Flow cytometry

The procedure for processing the spleen and isolating dendritic cells was followed as per the isolation protocol described above. Subsequently, the isolated cells were stained with a live/dead dye, which was diluted 1:1000 in PBS supplemented with a 1:250 dilution of Fc block for 10 minutes at room temperature. The cells were then stained with antibodies diluted according to the antibody table above in 1% FBS-containing PBS for 60 minutes on ice. After staining, the cells were washed twice and resuspended in 1% FBS-containing PBS for analysis on an Attune NxT Flow cytometer. The collected data was analyzed using FlowJo software version 10.4 and the resultant values were plotted using Graphpad Prism version 8.

### Correlative Focused Ion Beam Scanning Electron Microscopy (FIB-SEM)

RawKb.APOL7C::mCherry cells were seeded into a gridded glass ibidi μdish at 20% confluency. The culture medium was supplemented with doxycycline to a final concentration of 1 μg/ml and incubated for 4 hours at 37°C. After the first hour of doxycycline treatment, 50 μg of zymosan was added to the culture media and incubated for an additional 3 hours. The doxycycline and zymosan-containing culture medium was replaced with fresh medium before imaging at 37°C, 5% CO_2_ on Leica SP8 confocal microscope equipped with a HC PL APO 63×/1.40 Oil CS2. Following confocal imaging, the ibidi dish was washed twice with PBS to remove the culture medium and immediately fixed with 2% PFA + 2.5% glutaraldehyde in 0.2 M cacodylate buffer for 1 hour. The cells were then washed three times with 0.15 M cacodylate buffer and stained with 2% OsO_4_ in 1.5% potassium ferrocyanide-containing cacodylate buffer for 90 minutes. The ibidi dish was washed three times with 0.15 M cacodylate buffer and dehydrated with a series of ethanol concentrations (50%, 70%, 90%, 95%, and 100%) and pure acetone for 15 minutes each. The Durcupan resin was prepared by combining part A (11.4g), part B (10g), part C (0.3g), and part D (0.1g). For better infiltration, the resulting resin was mixed with acetone in ratios of 3:1, 1:1, and 1:3, and added sequentially to the ibidi dish after acetone dehydration for infiltration, with each step taking 30 minutes and a final overnight step at 4°C with 100% resin. On the following day, the overnight resin was replaced with fresh resin and polymerized in a 60°C oven for 48 hours.

The serial sectioning and imaging was performed on a dual-beam FIB-SEM Helios 5 CX (Thermo Fisher Scientific Inc.) equipped with a gallium liquid metal ion source. The resin blocks were mounted on an SEM stub using a conductive epoxy resin glue and with silver resin paste and then sputter-coated with carbon (Leica EM ACE600) to enhace sample conductivity before being loaded onto a multi-purpose holder in the SEM sample chamber. An overview low resolution back scattered electron (BSE) image of the entire block was acquired with the Everhart-Thornley detector to identify cells of interest (COI) by comparing with the confocal image. Once the COI was identified, the sample was positioned at the eucentric height and was prepared for the serial-sectioning and imaging (FIB-BSE) workflow. Briefly, a ∼1–2 µm thick protective layer of platinum was deposited on the top of the COI using a gas injection system (GIS) and trenches (∼60 µm (X) × 30 µm (Y) × 15 µm (Z)) were dug in front and along each side (∼ 10 µm wide) of the COI using the focused ion beam (FIB) to expose its block face. Once prepared, the COI was sequentially milled using the FIB and imaged using the SEM and BSE signals via an automated serial sectioning and imaging workflow (Auto Slice & View 4.2 (ASV) software package). SEM and BSE images were acquired using 2kV accelerating voltage, x mm working distance, and at a beam current of 0.69 nA with the Elstar in-lens (TLD) and the Elstar in-column (ICD) detectors, respectively, and typical dwell times of 3-5 μs. This allowed us to obtain high-resolution three-dimensional (3D) images (6-8 x 6-8 x 10 nm^3^ voxels) of the COI. The 3D images were then aligned and segmented manually using the TrakEM2 plugin in ImageJ (version 2.9.0/1.53t, build a33148d777) and registered and reconstructed in Imaris 10.0.0 (Oxford Instruments).

### Statistical analysis

GraphPad Prism software was used to perform all statistical analyses. Statistical significance between two samples was determined using an unpaired, two-tailed Welch’s Student’s *t* test, Kolmogorov-Smirnov test, or unpaired parametric Mann-Whitney tests as indicated in the figure legends. Data are displayed as mean ± standard deviation or ± standard error of the mean as indicated in the figure legend.

## Acknowledgements

We would like to thank Dr. Sergio Grinstein, Dr. Jayne Raper, Dr. Caetano Reis e Sousa and members of the Canton Lab for helpful discussions. We thank Dr. Xuejun Sun and Dr. Yuhuan Zhou for their help with the correlative FIB-SEM. We would also like to thank Dr. Paul Gordon for assistance with the bulk RNAseq and Dr. Karen Poon for assistance with flow cytometry and sorting.

## Author contributions

G.A.G., S.H. and J.C. conceived of the project, designed, and performed the experiments. J.C. wrote the manuscript. J.R., L.W., C.W., J.A.N., M.M., E.C., N.M., and M.B.C. designed and performed experiments. V.E. performed some of the microscopy experiments. N.R. ran the scRNAseq and performed the analysis. S.S. assisted with analysis of the scRNAseq. B.S. helped to run the *S. aureus* experiments. N.C.P. assisted with the characterization of immune compartments in the APOL7C knockout mice. J.B. helped design the scRNAseq experiment. D.J.M. helped design the immunization experiments. R.M.Y. helped to design some of the phagocytosis assays. This work was supported by a start-up grant from the University of Calgary to J.C., the VPR Catalyst Grant from the University of Calgary to J.C., an NSERC Discovery Grant to J.C., a CIHR Project Grant to J.C., and the Bhagirath Singh Early Career Prize in Infection and Immunity to J.C.

## Competing Interests

The authors declare no competing interests.

## Data and Code Availability

The bulk and single-cell RNA-seq will be deposited in the NCBI Genome Expression Omnibus and Sequence Read Archive once the paper is accepted for publication. Prior to this, it will be made available upon reasonable request. This paper does not report original code.

---

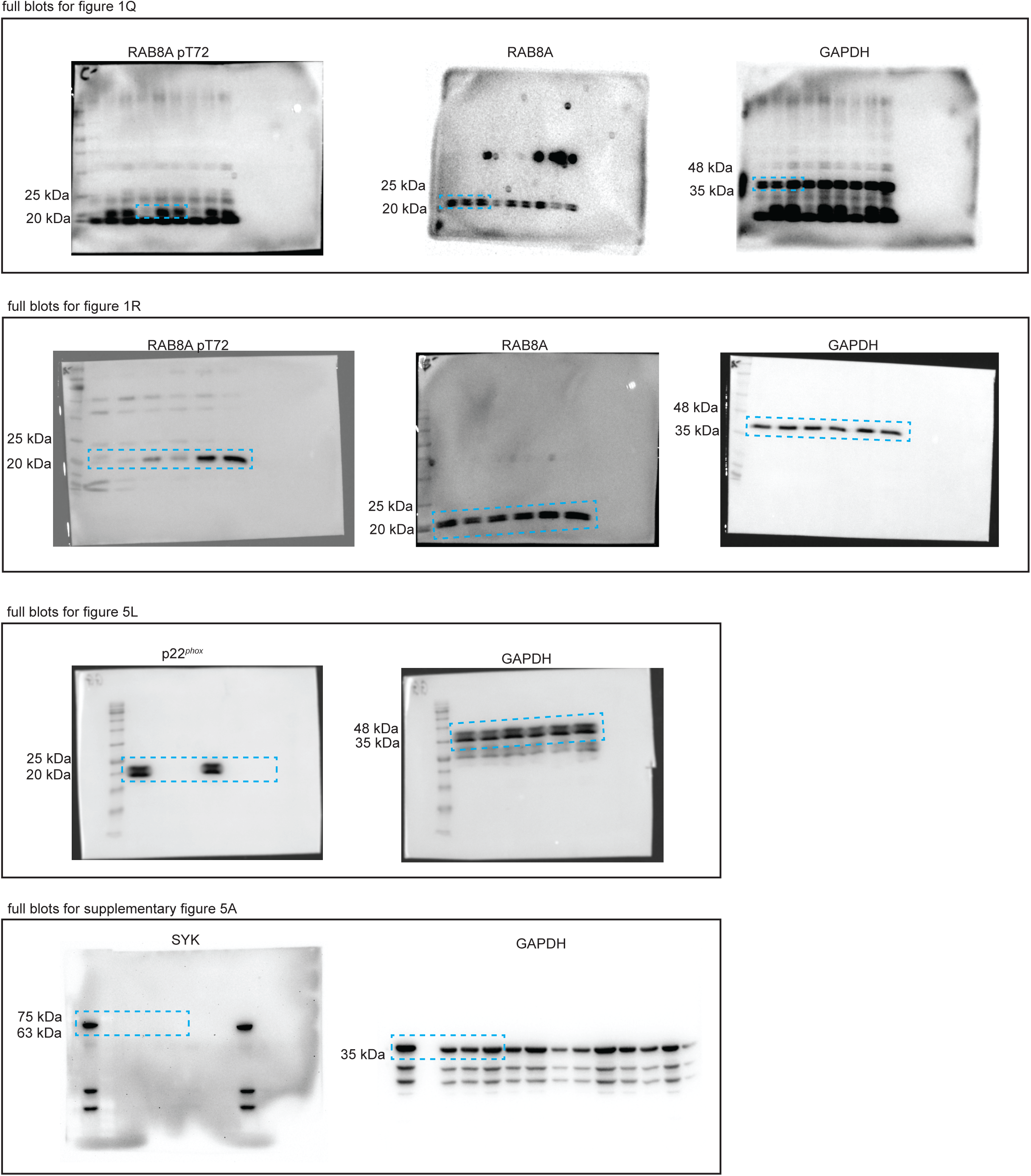

## Notes

### Competing Interest Statement

The authors have declared no competing interest.

### Summary of Updates

We have added a new experiment to Figure 4. We now include data for immunizations of Apol7c+/- and Apol7c-/- mice. We have also added a new supplementary figure (Supplementary figure 7) which show flow cytometric immune profiling of cells from the spleens of Apol7c+/- and Apol7c-/- mice.

